# Respiratory supercomplexes provide metabolic efficiency in zebrafish

**DOI:** 10.1101/818286

**Authors:** Carolina García-Poyatos, Sara Cogliati, Enrique Calvo, Pablo Hernansanz-Agustín, Sylviane Lagarrigue, Ricardo Magni, Marius Botos, Xavier Langa, Francesca Amati, Jesús Vázquez, Nadia Mercader, José Antonio Enriquez

**Author notes:** Correspondence should be addressed to: N. Mercader or J.A. Enríquez.

## Abstract

The oxidative phosphorylation (OXPHOS) system is a dynamic system in which the respiratory complexes coexist with super-assembled quaternary structures called supercomplexes (SCs). The physiological role of SCs is still disputed. Here we used zebrafish to study the relevance of respiratory SCs. We combined immunodetection analysis and deep data-independent proteomics to characterize these structures and found similar SCs to those described in mice, as well as novel SCs including III_2_+IV_2_, I+IV and I+III_2_+IV_2_. To study the physiological role of SCs, we generated two null allele zebrafish lines for supercomplex assembly factor 1 (SCAF1). SCAF1^-/-^ fish displayed altered OXPHOS activity due to the disrupted interaction of complex III and IV. SCAF1^-/-^ fish were smaller in size, and showed abnormal fat deposition and decreased female fertility. These physiological phenotypes were rescued by doubling the food supply, which correlated with improved bioenergetics and alterations in the metabolic gene expression program. These results reveal that SC assembly by SCAF1 modulates OXPHOS efficiency and allows for the optimization of metabolic resources.

---

In the last two years, the focus of investigation on the structure of the mitochondrial electron transport chain (ETC) has shifted from the dispute over the existence of supercomplexes (SCs) to their putative functional role. In mammals, the best understood mechanism of respiratory complex super-assembly is the interaction between complexes III (CIII) and IV (CIV) mediated by supercomplex assembly factor 1 (SCAF1/COX7A2L)^1^. The carboxy-terminus of SCAF1 is very similar to that of the CIV subunit COX7A2 and replaces it in the subset of CIV molecules that super-assemble with CIII^2^. After some initial doubts^3^, which were later dispelled^4, 5^, the role of SCAF1 in the super-assembly of CIII and CIV is now generally accepted^2^. The process of super-assembly between CI and CIII and CI and CIV to form the respirasome is unknown, but the proposed existence of I+IV SCs^6^ suggests that CI-CIII and CI-CIV super-assembly might occur independent from CIII and CIV assembly^7, 8^. So far, the interaction between CI and CIV has been mostly studied in SCs containing CI, CIII and CIV (also named respirasomes). Several forms of respirasomes (I+III2+IV) migrate closely together in blue native gel electrophoresis (BNGE), although the reason for their different apparent molecular weights remains unknown. Even though SCAF1 loss-of-function abolishes the interaction between CIII and CIV, the current consensus is that respirasome formation is not completely disrupted in the absence of functional SCAF1. However, the stability and the variety of respirasomes is strongly reduced in the absence of functional SCAF1^1, 2, 4^.

The super-assembly between CI and CIII was proposed to allow partitioning of coenzyme Q (CoQ) into two communicated functional pools: one trapped in SCs and the other free within the inner mitochondrial membrane ^9^. The super-assembly between CIII and CIV allows the control of available CIV through compartmentalization. Both functions optimize the metabolic flux, preventing an electron traffic jam^1^ and minimizing reactive oxygen species (ROS) production^10^ while maintaining an efficient energy production^9^. This model was, however, contested by studies performed in fragmented sub-mitochondrial particles generated by disruption of mitochondrial membranes with detergents^11^. These studies concluded that CoQ pools are continuously intermixed at a rate that rules out the possibility of preferential use of CoQ within SCs. Accordingly, these studies defended the notion that the super-assembly between CI and CIII in the form of I+III_2_ or I+III_2_+IV would lack any bioenergetic role^5^. A very recent publication analyzing isolated supercomplex I+III_2_ also support the model were partitioning of CoQ SC: I+III_2_ have functional implications in the oxidation of NADH^12^.

*In vivo*, the functional role of SCs has been supported by the positive correlation of the proportion of respirasomes (I+III_2_+IV) with exercise^13^, and with lower mitochondrial ROS production^14–16^. Interestingly, some common mouse strains, such as C57BL/6, have a non-functional SCAF1 protein (SCAF1^111^) lacking two amino acids as compared with functional SCAF1 (SCAF1^113^). SCAF1^111^ mice do not assemble III_2_+IV and the formation of respirasomes is affected^1, 2^. No specific phenotype has yet been ascribed to SCAF1^111^ mouse strains, and this has contributed to the arguments against a physiological role for the super-assembly of CIII and CIV. Furthermore, a recent study performed in cultured cells suggested that the ablation of SCAF1 has no bioenergetic relevance^17^, a conclusion that was refuted by the demonstration that SCAF1 was critical for the cellular metabolic adaptation to the increase in energy demands associated with endoplasmic reticulum stress and nutrient starvation^18^. Controversy over the bioenergetic role of CIII and CIV super-assembly thus remains, and a physiological role has not been yet described. Here, we studied respiratory SCs in zebrafish (*Danio rerio*), and evaluated the super organization and plasticity of the oxidative phosphorylation (OXPHOS) system. We identified SCs as those previously described in mice, such as I+III, III_2_+IV and I+III_2_+IV, but also novel SCs, including III_2_+IV_2_, I+IV and I+III_2_+IV_2_. As suggested for mammals, super-assembly dynamics are dependent on the metabolic status in this vertebrate model organism. With the aim of studying the bioenergetic and physiological consequences of impaired super-assembly, we generated SCAF1 null mutant zebrafish lines. SCAF1^-/-^ fish showed a disrupted interaction of CIII and CIV, and presented with a prominent phenotype, including a smaller size despite having higher body fat accumulation. In addition, males showed a reduced swimming capacity, and females decreased fertility. Bioenergetics analysis revealed that SCAF1 ablation promotes an inefficient OXPHOS capacity due to the disruption of the compartmentalization of CIV. Strikingly, phenotypic alterations in SCAF1^-/-^ animals are fully corrected by doubling the food supply but not by changing the regime to a high-fat diet. The phenotypic rescue occurred in the absence of a recovery of OXPHOS super-assembly and correlated with alterations in the metabolic gene expression program.

Overall, these results confirm a role for SCs in the efficiency of the respiratory chain in a vertebrate animal model and reveal that SCs provide an advantage in the optimization of metabolic resources.

## RESULTS

### OXPHOS super-assembly in zebrafish

To re-evaluate whether the disruption of SCs formation has any impact at the organismal level, here we used the zebrafish animal model. OXPHOS genes have been reported to be highly conserved along evolution^19^ and the interest in the use of zebrafish in metabolism and mitochondrial biology is increasing^20^. Nonetheless, the super-assembly of respiratory complexes in this model has not been examined. Thus, we first characterized the pattern of respiratory complex super-assembly in the zebrafish using blue native gel electrophoresis (BNGE). Purified zebrafish skeletal muscle mitochondria were run in parallel with mitochondria from mouse skeletal muscle of either functional SCAF1^113^ (CD1) or non-functional SCAF1^111^ (C57BL/6J) mice (Fig. 1a-d). As expected, the III_2_+IV SC was absent in SCAF1^111^ mitochondria and the abundance of the respirasome (I+III_2_+IV) was very low. The BNGE pattern of zebrafish CI and CIII was similar to that of SCAF1^113^ mice, where free I, III_2_ and I+III_2_ could be easily identified (Fig. 1a and Fig. S1a, c to see the split channels of Fig 1a). The migration pattern of zebrafish CIV paralleled that of mice, and revealed the presence of IV_2_ and III_2_+IV in zebrafish. It also confirmed the presence of the respirasome, which migrated faster than that of mice (Fig. 1b and Fig. S1a-c to see the split channels of Fig 1b). We also noted some conspicuous differences between the pattern in zebrafish and the classical mouse pattern; specifically, the co-migration of CIII and CIV just above free CI (Fig 1b, Fig. S1c), as well as the co-migration of CI and CIV just below I+III_2_, which is compatible with the interaction of CI and CIV. The CI and CIV interaction was barely detectable by immunodetection (Fig. S1a, c), but could be clearly revealed by CI (Fig. 1c) and CIV (Fig. 1d) in-gel activity, which also confirmed the other observations based on immunodetection.

**Figure 1.**
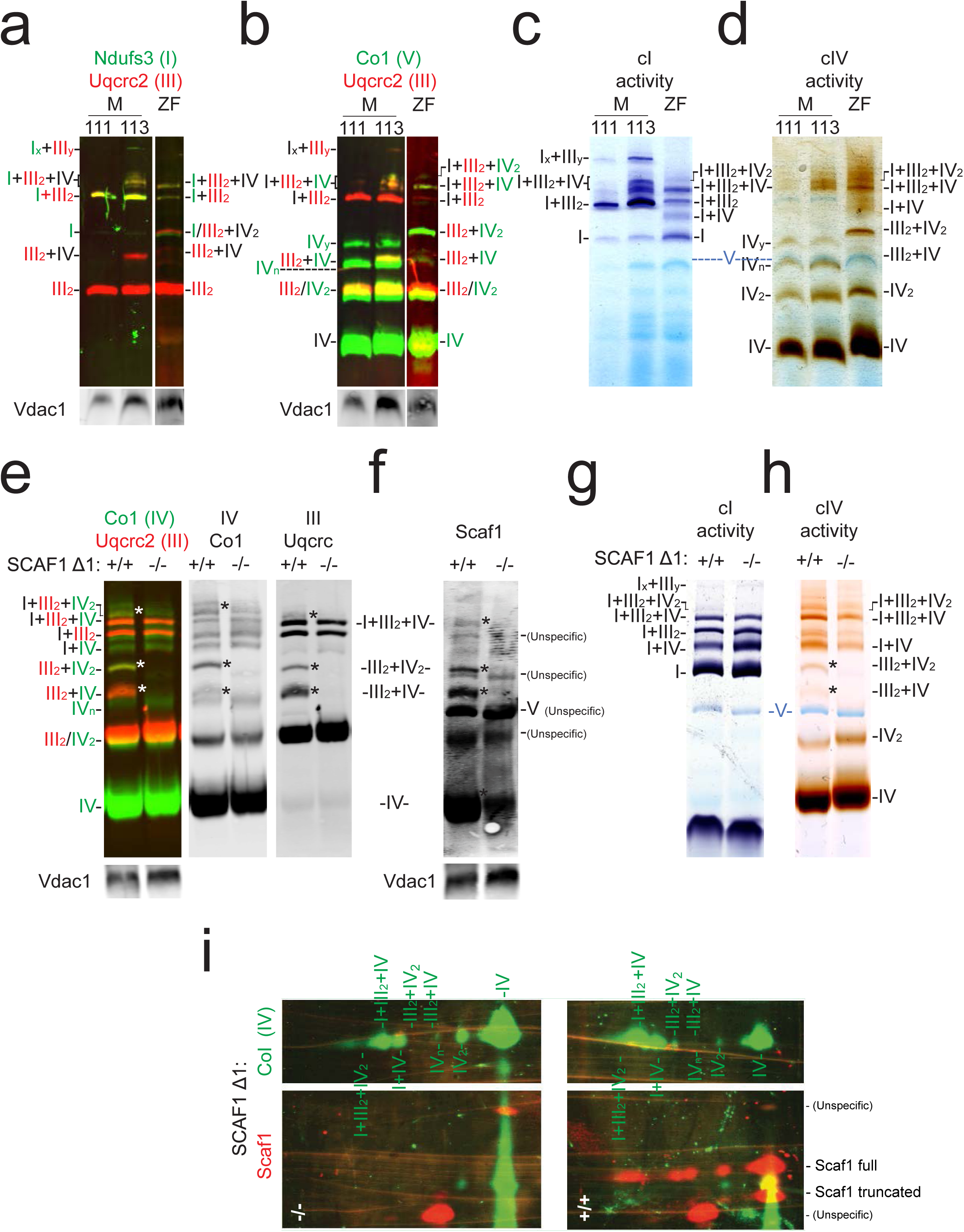
OXPHOS super-assembly in zebrafish. **a-d**, Blue native gel electrophoresis (BNGE) of mouse (M) C57BL/6J (111), CD1 (113) and zebrafish (ZF) skeletal muscle digitonin-solubilized mitochondria. (**a, b**) Immunodetection of the indicated proteins after BNGE, (**c**) in-gel activity for CI and (**d**) CIV (shown is a representative gel from two technical and two biological replicates). **e-h**, BNGE of whole-body zebrafish digitonin-solubilized mitochondria of SCAF1^Δ1/Δ1^ and its respective SCAF1^+/+^ counterpart. (**e, f**) Immunodetection of the indicated proteins, (**g**) in-gel activity of CI and (**h**) CIV (representative gel from two technical and three biological replicates). **i**, 2D BNGE/SDS electrophoresis: 1^st^ dimension with digitonin (Dig) and 2^nd^ dimension with SDS, followed by immunoblotting with the indicated antibodies to identify the proteins detected by the commercial anti-SCAF1 antibody.

The novel band containing CIII and CIV is compatible with the presence of a dimer of CIII with two molecules of CIV. Two alternative structural arrangements could generate this band – the interaction of two dimers, III_2_ and IV_2_ (III_2_+IV_2_), or the interaction of one dimer of III with two monomers of IV (IV+III_2_+IV or 2IV+III_2_). To distinguish between these possibilities, we performed two-dimensional (2D)-BNGE using digitonin as a detergent in the first dimension to preserve the integrity of SCs, and n-dodecyl-β-D-maltoside (DDM) in the second dimension to disaggregate SCs while substantially preserving the integrity of the complexes^21^ (Fig. S1d, e). This analysis allowed us to differentiate between co-migration and true DDM-sensitive interactions, and to discern whether CIV was in a monomeric or dimeric form. Although DDM partially disassembled IV_2_ into its monomers, we were still able to detect a considerable proportion of IV_2_. Indeed, we found a dimer of CIV (IV_2_) present in the novel high molecular weight band containing CIII and CIV, indicating that this co-migration is due to the physical interaction between III_2_ and IV_2_ (Fig. S1d-e). In addition, 2D-BNGE analysis demonstrated that CI and CIV co-migration was due to a physical interaction between them in the form of I+IV SC, which could be disrupted by DDM (Fig. S1d-e). Interestingly, this assay also revealed the presence of CIV dimers in high molecular weight structures associated with CI and CIII (putative I+III_2_+IV_2_) or with CI alone (putative I+IV_2_) (Fig. S1d-e).

These observations were confirmed in mitochondria purified from whole zebrafish (Fig. S2a-e). Of relevance, the pattern of zebrafish BNGE bands was stable between 4 and 10 g/g proportion of digitonin whereas 1 g/g proportion was unable to sufficiently solubilize the mitochondrial membranes (Fig. S2f).

To evaluate whether the proposed role of respiratory complex super-assembly plasticity during the adaptation to variations in metabolic resources^1, 9^ applies to zebrafish, we fed adult zebrafish for 6 weeks with a low proteins/low fats (LP/LF) diet, which leads to malnutrition (Fig. S2g-i). LP/LF-fed fish showed a decrease in weight, which was accompanied by an increase of I+III_2_ and I+III_2_+IV_1-2_ SCs and a decrease of III_2_+IV_1-2_. (Fig. S2i). These results confirm that the organization of the ETC is modulated in response to metabolic changes also in zebrafish.

Given the similarities with mice, its dynamic plasticity according to the metabolic status, and the possibility to study novel SCs such as I+IV and those containing IV_2_, we conclude that the zebrafish is a valuable model to study SC assembly and function.

### Role of SCAF1 in zebrafish respiratory complexes super-assembly

To assess structural and physiological effects of alterations in complex super-assembly we generated a zebrafish SCAF1 null mutant model. Zebrafish SCAF1, also known as *cox7a2l* and *cox7a3*, is highly conserved compared with mammals (Fig. S3a). We generated two independent SCAF1 zebrafish knockout lines (Fig. S3a) using CRISPR/Cas9 technology (Δ1 and Δ2; Fig. S3b-c). We introduced premature STOP codons after amino acids 43 and 51, respectively, which, according to sequence information, leads to a short nonfunctional protein (Fig. S3d-e). As expected, we could not detect SCAF1 protein by western blotting of SCAF1^Δ1^ and SCAF1^Δ2^ homozygous fish (Fig. S3f). We next extracted mitochondria from whole SCAF1^+/+^ and SCAF1^-/-^ fish and compared SC formation by BNGE (Fig. 1e-h). The lack of SCAF1 completely eliminated the two bands of SCs containing CIII and CIV (III_2_+IV, III_2_+IV_2_). It also strongly diminished the I+III_2_+IV_2_ band, whereas the I+III_2_+IV SC was only slightly decreased in intensity. Immunodetection of CIII, CIV and SCAF1 (Fig. 1e-f) as well as CI and CIV in-gel activity assays (Fig. 1g-h) confirmed these observations.

SCAF1 was immunodetected in BNGE in a greater number of bands than those reported in mammals, from which three completely disappeared in SCAF1^-/-^ samples (Fig. 1f). We found that this is due to the discontinuation of the commercial SCAF1 antibody generated against a SCAF1 specific peptide, and its substitution by another generated against the entire protein. The new antibody caused the appearance of strong non-specific immunoreactive bands. To distinguish between SCAF1-specific and non-specific immunodetected signals, we performed 2D-BNGE/SDS-PAGE electrophoresis of SCAF1^+/+^ and SCAF1^-/-^ samples (Fig. 1i). This analysis revealed that the antibody recognized four proteins of different molecular weight. Two of them remained present in the SCAF1^-/-^ sample, indicating that they correspond to spurious signals. From the two specific signals that disappeared in SCAF1^-/-^ samples, one matched with the expected size of SCAF1 and migrated in 6 different spots corresponding to complexes IV, IV_2_, III_2_+IV, III_2_+IV_2_, I+III_2_+IV and I+III_2_+IV_2_. Unexpectedly, a second specific protein with lower molecular weight was present in two spots (IV, IV_2_), suggesting the presence of a shorter form of SCAF1 (Fig 1i).

To further study the effect of SCAF1 loss-of-function on SC assembly, we performed high-throughput Blue-DiS-based proteomics of the entire BNGE run of SCAF1^+/+^ and SCAF1^-/-^ samples. This allowed the detection of around 9,000 UNIPROT entries (Supplementary Data Sheet 1). In the currently annotated zebrafish proteome most of the UNIPROT entries are unreviewed, a large proportion is referred to as fragments, and a great number of entries correspond to unidentified proteins (Supplementary Data Sheet 1). Therefore, some entries may correspond to the same gene and would not accurately report the number of different proteins within the gel. Proteins were detected at a very variable concentration covering 6 orders of magnitude, from which mitochondrial proteins (Fig. S4a), and in particular the inner membrane proteins, were the most abundant (Fig. S4a-b). Proteomic analysis confirmed the pattern of OXPHOS complexes obtained by immune analysis and in-gel activity (Fig. 2a and Fig. S4c-d). It also revealed the high reserve of the CI N-module retaining its NADH dehydrogenase activity (Fig. 1g and Fig. S4c-d). This accumulation was paralleled by a significant amount of CI lacking NADH activity that co-migrated with SC III_2_+IV (Fig. 1g and Fig. S4c-d). Of note, SCAF1-derived peptides produced a negligible signal in the SCAF1^-/-^ sample (Fig. S4c-d).

**Figure 2.**
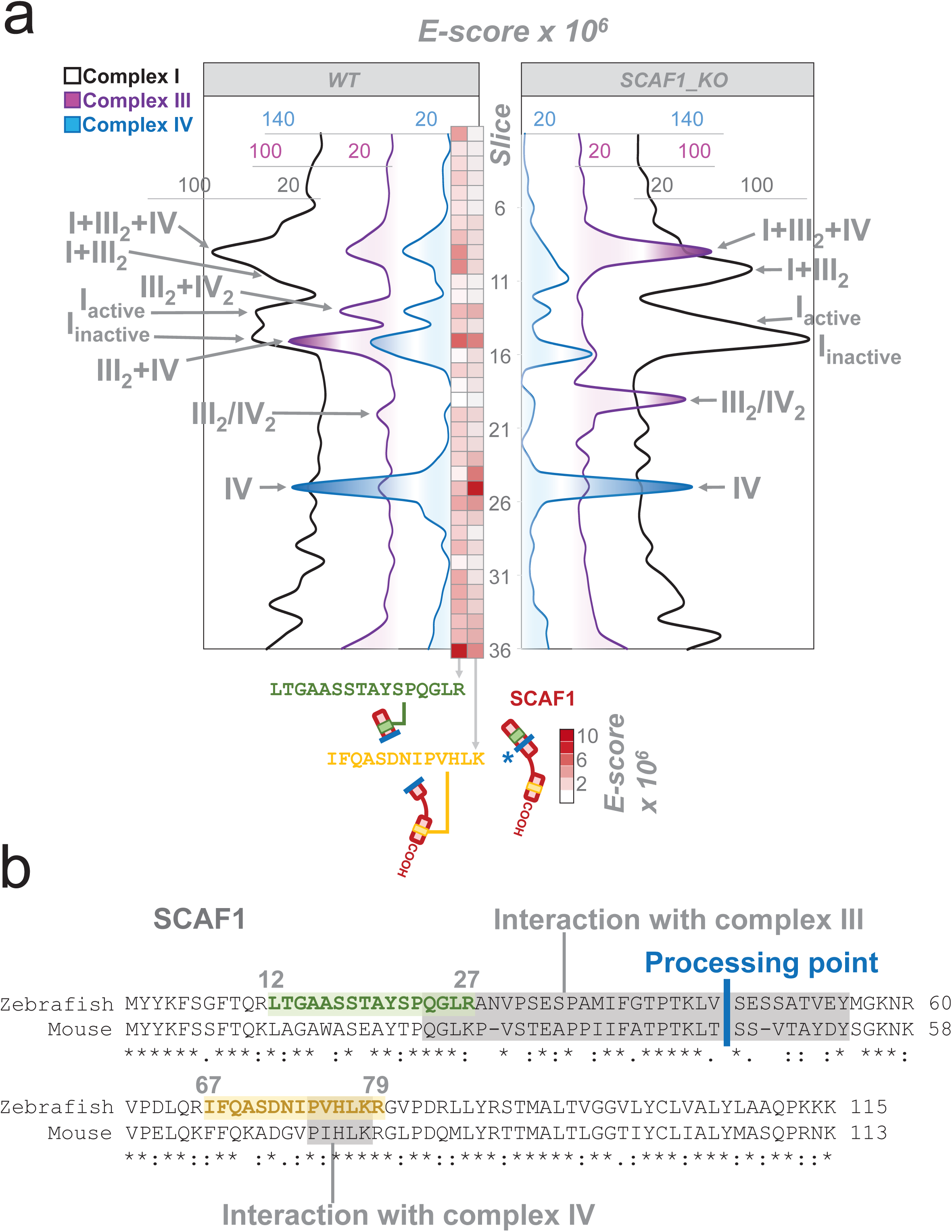
Blue-DiS proteomics of SCAF1^+/+^ and SCAF1^-/-^. **a**, Quantitative data-independent scanning (DiS) mass spectrometry protein profiles for CI, CIII and CIV. Vertical numbers indicate the BNGE gel slices. Left and right profiles correspond to SCAF1+/+ and SCAF1^Δ1/Δ1^ animals, respectively. Red heatmap corresponds to the E-score from two proteotypic SCAF1-derived tryptic peptides spanning sequences ascribed to the CIII interacting site (in green) and to the CIV interacting site (in yellow). Thick blue line, marked with an asterisk, indicates the putative proteolytic site in SCAF1. **b**, Sequence alignment of SCAF1 protein in mouse and zebrafish. Structural and functional regions previously described in mouse are indicated in shaded grey boxes. Thick blue line indicates the proteolytic processing site in the mouse sequence.

The proteomic analysis also allowed a more quantitative estimation of the changes induced by SCAF1 ablation (Fig. S5a). It confirmed that the absence of functional SCAF1 impaired the super-assembly of CIII with CIV (Fig. S5a), and showed that both CIII and CIV completely disappeared from III_2_+IV_2_ and III_2_+IV bands, with the concomitant increase in III_2_ and IV. Conversely, the electrophoretic mobility of the active and inactive forms of CI were unaffected, demonstrating that they co-migrate but do not interact (Fig. S5a, b). Nevertheless, we noticed a significant quantitative shift from the active to the inactive form of CI and a parallel increase in the amount of the free CI N-module. Proteomic analysis also confirmed that the different forms of respirasomes were still present in the SCAF1^-/-^ samples. However, it revealed quantitative differences when comparing SCAF1^-/-^ and controls, with a reduction in I+III_2_+IV and in the putative I_2_+III_2_+IV_2,_ as well as an increase in I+III_2_, the putative I_2_+III_2_ and the amount of I+IV (Fig. S5a, b).

Proteomic analysis confirmed that the smaller protein co-migrating with IV and IV_2_ was indeed a version of SCAF1 missing its amino terminus (Fig. 2a). By monitoring the presence of two proteotypic peptides for zebrafish SCAF1 located either at the amino- (CIII-interacting domain) or carboxy- (CIV-interacting domain) portions of the protein (Fig. 2b), we were able to determine whether SCAF1 was present in full or truncated forms. We found that the proportion of the two peptides was similar (slice 15, red-scaled heatmap in Fig. 2a), demonstrating the presence of full-length SCAF1 only in the bands where CIII and CIV interact. By contrast, when SCAF1 was found in free CIV (slice 25, red-scaled heatmap in Fig. 2a), the abundance of the amino-peptide was severely decreased and only the carboxy-peptide was detected in significant amounts. We speculate that the short SCAF1 form might derive from the proteolytic cleavage of full-length SCAF1 in a position located in the sequence of interaction with CIII (processing point Fig 2b), disrupting the interaction between CIII and CIV and giving rise to the free complexes. This interpretation is consistent with the fact that the SCAF1 amino terminus appeared at the bottom of the gel (Fig 2a).

### Physiological consequences of the disrupted supercomplex assembly in SCAF1^-/-^ zebrafish

Interestingly, the ablation of SCAF1 caused a prominent phenotype in zebrafish (Fig. 3). Both males and females were significantly shorter (Fig. 3a-d) and females weighed significantly less (Fig. 3e-f). Moreover, SCAF1^-/-^ animals showed poorer swimming performance, although the differences were significant only in males (Fig. 3g-h).

**Figure 3.**
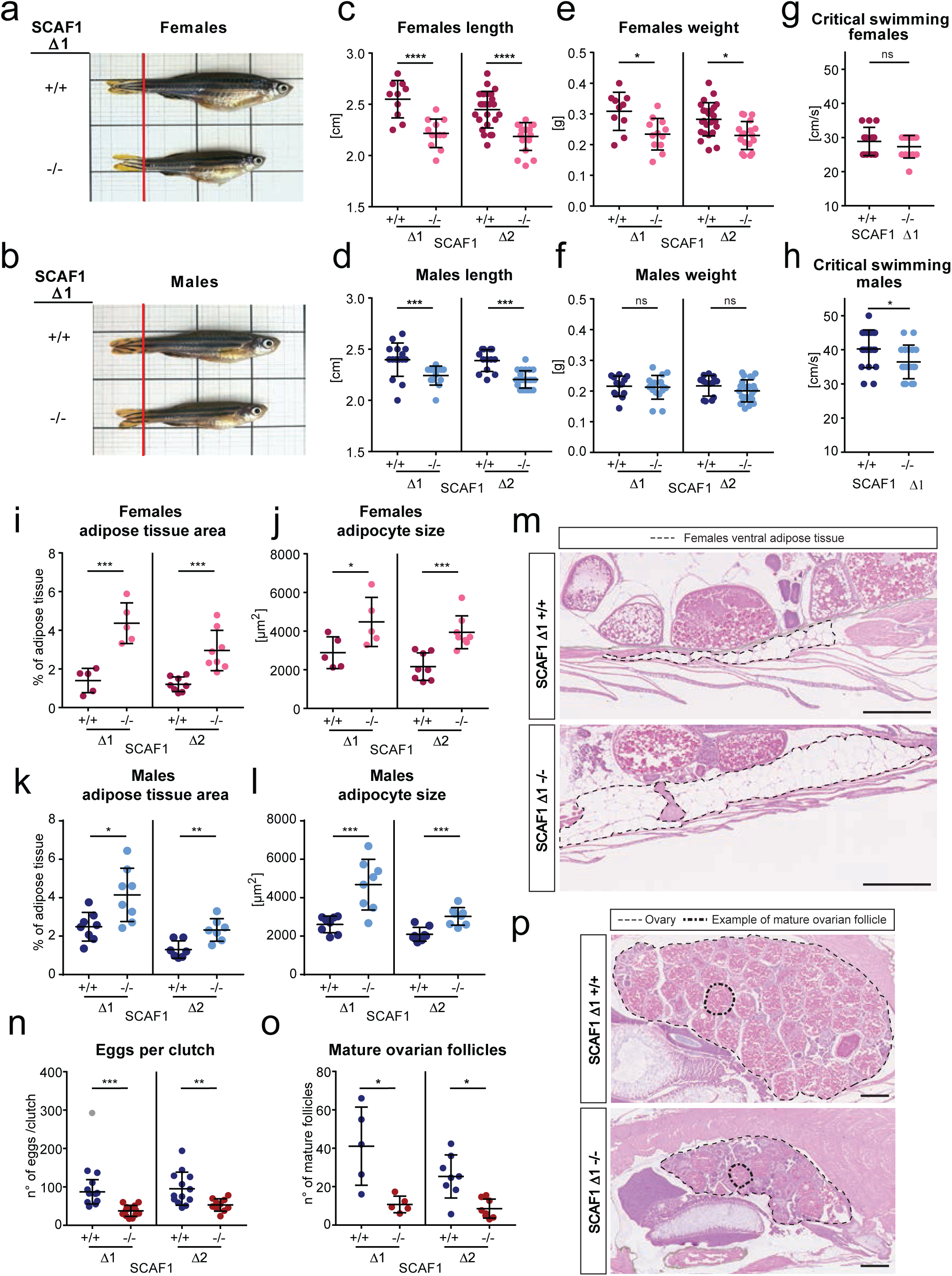
Phenotype consequences of SCAF1 loss-of-function. **a, b,** Representative images from SCAF1^+/+^ and SCAF1^-/-^ (**a**) female (**b**) male adult zebrafish. **c-f,** Size of SCAF1^Δ1/ Δ1^ and SCAF1^Δ2/ Δ2^ fish in comparison with their respective SCAF1^+/+^ lines, (**c**) length and (**e**) weight of females (Δ1 +/+ n=10, Δ1 -/- n=12, Δ2 +/+ n=24, Δ2 -/- n=18); (**d**) length and (**f**) weight of males (Δ1 +/+ n=16, Δ1 -/- n=13, Δ2 +/+ n=13, Δ2 -/- n=23). **g,h,** Swimming performance of SCAF1^Δ1/ Δ1^ fish in comparison with their respective SCAF1^+/+^ fish. Critical swimming speed of (**g**) females (+/+ n=13, -/- n= 13) and (**h**) males (+/+ n= 18, -/- n=17). **i-m,** Adipose tissue measurements on hematoxylin-eosin (H&E)- stained adult zebrafish sagittal sections. (**i, k**) Adipose tissue area per total section area (average of 3 sections/biological replicate) and (**j, l**) adipocyte size (average of 20–30 adipocytes of ventral adipose tissue per biological replicate) of females (**i, j**) (Δ1 n=8, Δ2 n=8, same number of animals for homozygous mutants and controls) and males (**k, l**)(Δ1 n=8, Δ2 n=7, same number of animals for homozygous mutants and controls). **m,** Representative images of ventral fat deposits in females (dotted lines). n-p, Effect of SCAF1 loss of function on female fertility. **n**, Number of eggs per clutch (Δ1 +/+ n=12, Δ1 -/- n=13, Δ2 +/+ n=13, Δ2 -/- n=10). **o**, Quantification of mature ovary follicles per ovary section (average of three sections/biological replicate) (Δ1 n=5, Δ2 n=8; same number of animals for homozygous SCAF1^+/+^ and SCAF1^-/-^). **p**, Representative images of H&E-stained ovaries. One-way ANOVA in **c-f**, **i-m** and unpaired t-test in **g,h**. Outliers are shown in grey and were not considered for the statistical analysis. Data are represented as mean ± SD. * p<0.05, ** p<0.01, *** p<0.001, **** p<0.0001. Scale bars = 500 μm.

We further observed that SCAF1^-/-^ animals accumulated more adipose tissue and showed increased adipocyte cell size (Fig. 3i-m), indicating an altered metabolism. In addition, SCAF1^-/-^ females showed a decrease in fertility, with a lower number of eggs per clutch (Fig. 3n), possibly due to delayed oocyte maturation (Fig. 3o-p). These phenotypes, as well as the loss of super-assembly between CIII and CIV, were observed in both SCAF1^-/-^ zebrafish lines and were not observed in SCAF1^+/-^ fish (Fig. S7). In sum, SCAF1^-/-^ fish show phenotypes resembling malnourishment (reduced growth, compromised exercise capacity and reproduction efficiency) when they are fed in equal conditions as SCAF1^+/+^ fish^22, 23^. Therefore, SCAF1 loss-of-function impairs the proper energy conversion from nutrients, storing them in abnormally high amounts at the adipose tissue instead of using them for energy production.

### Mitochondrial consequences of the disrupted supercomplex assembly in SCAF1^-/-^ zebrafish

To determine whether the observed phenotypes could be directly related to OXPHOS function, we first used transmission electron microscopy (TEM) to establish whether the absence of SCAF1 induced any ultrastructural alterations in zebrafish mitochondria (Fig. 4a). We observed an increase in the mitochondrial area in SCAF1^-/-^ cardiomyocytes of heart sections (Fig. 4b), which displayed more fragmented organelles (Fig. 4c) and wider cristae (Fig. 4d). Increased cristae lumen width has previously been associated with the reduction of super-assembly of respiratory complexes in models of cristae junction disruption^24, 25^. No differences in mitochondria content were noted, as measured by the ratio of mitochondrial DNA to nuclear DNA (Fig 4e-h) and the enzymatic activity of citrate synthase (Fig. S6a-b), both in heart and skeletal muscle (Fig 4e-h). Thus, the increase of mitochondria area could be due to the increase in cristae lumen width and or to cell-type specificities, as TEM analysis was performed specifically in cardiomyocytes.

**Figure 4.**
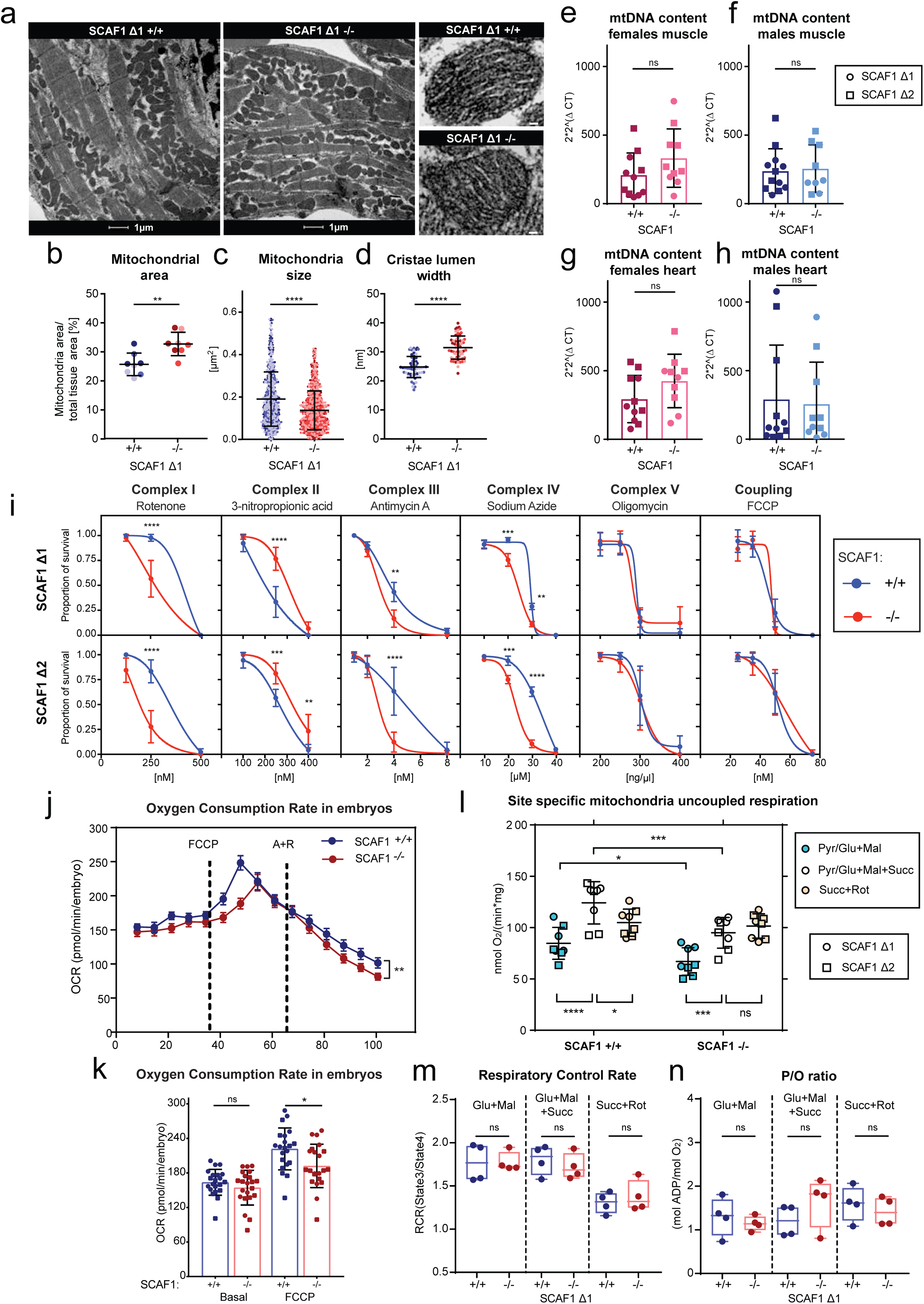
SCAF1 loss-of-function leads to alterations in mitochondrial structure and performance. **a-d**, Transmission electron microscopy image of cardiac muscle from SCAF1^Δ1/Δ1^ (n=3) and SCAF1^+/+^ fish (n=3). **a**, Representative images showing mitochondria. **b**, Mitochondria area per tissue area of cardiomyocytes (2–3 images per biological sample). **c**, Mitochondria size (100– 150 mitochondria per biological sample). **d**, Cristae lumen width (average of three cristae per mitochondria, 20 mitochondria per biological sample). Different biological replicates are represented with different color tones. **e-h**, Mitochondrial DNA copy number per nuclear copy number in muscle (**e, f**) and hearts (**g, h**), in females (**e, g**) and males (**f, h**) (Δ1 n=6, Δ2 n=6 and same number for their respective controls). **i**, Survival curve of 4 days post-fertilization embryos treated with different concentrations of the indicated OXPHOS inhibitors (three experimental replicates per biological replicate and three biological replicates). **j,k**, Oxygen consumption of 48 hours post-fertilization embryos using the XFe24 Seahorse analyzer, (**j**) Representative oxygen consumption rate (OCR) profile along time and (**k**) maximum OCR (Δ1 n=11, Δ2 n=11, n=10, n=11 respectively for their controls). **l**, Maximum uncoupled (FCCP) OCR in isolated mitochondria from adult fish (male and females Δ1 n=4 and Δ2 n=4, and same number for their respective controls) with the indicated site I [pyruvate (Pyr), glutamate (Glu), malate (Mal)] or site II [succinate (Succ)] substrates. **m, n**, Respiratory control rate (State 3/State4) (**m**) and P/O ratio (**n**) in isolated mitochondria from adult fish (male and females Δ1 n=4, and same number for their respective controls) with the indicated substrates. **b-h, j, k, m, n** Unpaired t-test, **i,** two-way ANOVA, Sidak’s multiple comparison test, **l,** RM two-way ANOVA, post-hoc Fisher’s LSD test. ns p>0.05, * p<0.05, ** p<0.01, *** p<0.001, **** p<0.0001. Data are represented as mean ± SD. Scale bars = large image 1 µm, small image 50 nm.

To further assess if OXPHOS function was affected, we determined the susceptibility of 4 days post-fertilization (dpf) SCAF1^+/+^ and SCAF1^-/-^ larvae to pharmacological inhibitors of the respiratory complexes I, II, III and IV, as well as inhibitors of H^+^-ATPase (CV) and mitochondrial coupling (Fig. 4i)^26, 27^. Notably, SCAF1^-/-^ embryos were significantly more sensitive than SCAF1^+/+^ embryos to the inhibition of CI, CIII and CIV, which participate in the formation of SCs, but were more resistant to the inhibition of CII (Fig. 4i), which does not super-assemble. In addition, the inhibition of CV (ATP synthesis) and the coupling between the ETC and ATP synthesis with FCCP were both insensitive to the presence of SCAF1 (Fig. 4i).

To gain direct insight on the impact of the SCAF1 ablation for mitochondrial respiration, we estimated the oxygen consumption capacity (OCR) of live mutant and wild-type zebrafish larvae using the Seahorse XFe24 analyzer^28^. Whereas the ablation of SCAF1 had no impact on the basal respiration (Fig. 4j-k), maximum oxygen consumption was significantly lower in zebrafish larvae lacking SCAF1. Notably, mitochondrial respiration of SCAF1^-/-^ larvae was more sensitive to antimycin and rotenone (Fig. 4j), providing a plausible explanation for the higher lethality of these drugs on SCAF1^-/-^ larvae (Fig. 4i).

The impact of SCAF1 ablation on mitochondrial respiration performance was also assessed in isolated mitochondria from adult zebrafish (Fig. 4l-n). Both respiratory control rate (RCR) and phosphate/oxygen ratio (P/O ratio) were unaffected by the loss of SCAF1 (Fig. 4m-n). However, we again observed that maximum respiration was significantly lower in SCAF1^-/-^ zebrafish mitochondria than in wild-type counterparts (Fig. 4l). We measured oxygen consumption in the presence of site I (pyruvate/glutamate and malate) or site II (succinate) substrates, finding that the decrease in maximum respiration was due to a decrease in site I substrate respiration, with site II respiration unaffected (Fig. 4l). Additionally, we observed that the respiration of wild-type zebrafish mitochondria with the combination of site I and site II substrates was higher than that obtained with only site I or site II substrates. This phenomenon was described previously in mouse^1^ and bovine^29^ mitochondria. Of note, site II substrates alone allowed a respiration level similar to that with combined site I and site II substrates in zebrafish SCAF1^-/-^ mitochondria, reproducing our observations in SCAF1-deficient mouse mitochondria^1^. Moreover, we found no differences in individual enzymatic activity of respiratory complexes in SCAF1^-/-^ fish (Fig. S6c-f). This strongly suggests that differences in OXPHOS performance and resistance to OXPHOS inhibitors are due to a different super-assembly and are not a consequence of changes in individual complex activities. In sum, these results are in agreement with the ***plasticity model*** of organization of the mitochondrial ETC, which proposes that super-assembly is not required for basal function of the ETC, but is required for organizing the flux of electrons and adapting to the differential balance of electrons arriving from different carbon sources^1^. These analyses demonstrate that the loss of SCAF1 has a direct functional impact on both OXPHOS performance and mitochondria structure. It also reveals a physiological role of a correct OXPHOS assembly at the organismal level.

### Diet determines SCAF1^-/-^ physiological phenotypes

The overall phenotype of SCAF1^-/-^ animals is reminiscent of a response to a mild decrease in mitochondria performance. Contrary to the *ad libitum* method of feeding mice, zebrafish receive food in defined doses. Since we had observed that changes in food supply affected OXPHOS levels (Fig. S2g-i), we wondered whether increasing food availability could ameliorate the phenotype of SCAF1^-/-^ animals. We fed SCAF1^+/+^ and SCAF1^-/-^ fish with the double amount of food, which was distributed in more doses per day to ensure complete intake. Strikingly, this treatment was sufficient to eliminate the differences in growth both in females (Fig. 5a-c) and in males (Fig. S8) after only 4 weeks. As expected, SCAF1^+/+^ animals fed with double diet showed a prominent increase in adipose tissue area and adipocyte size, but these changes were not as evident in SCAF1^-/-^ animals (Fig. 5d-f). The increase in food also rescued female fertility in SCAF1^-/-^ fish (Fig 5g-i).

**Figure 5.**
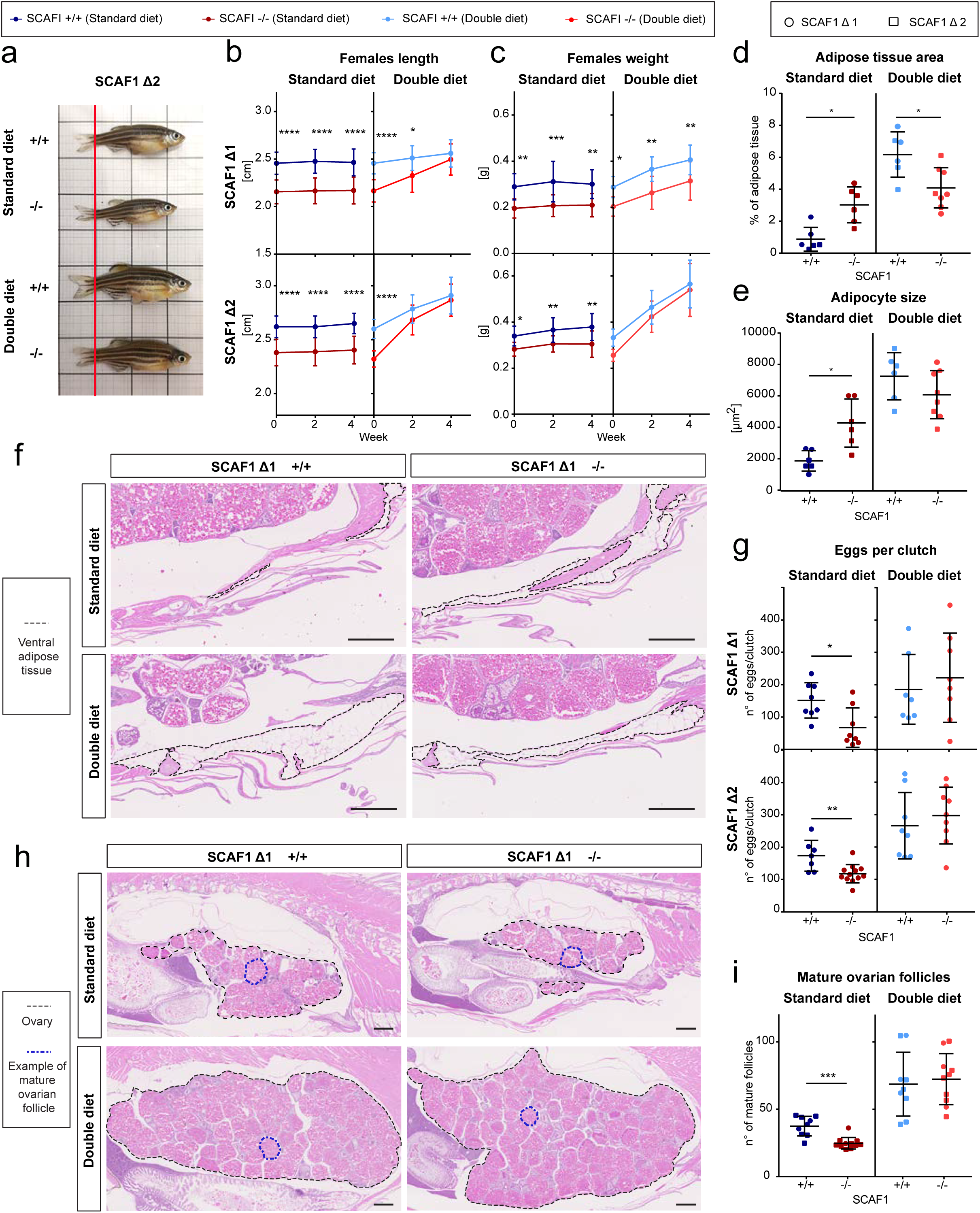
Diet-induced recovery of SCAF1^-/-^ phenotypes. Data from females. **a**, Representative images of SCAF1^-/-^ and SCAF1^+/+^ fish fed with the indicated diets. **b**, Changes in length and **c**, weight over time (Δ1 +/+ n=10, Δ1 -/- n=10, Δ2 +/+ n=10, Δ2 -/- n=12-13. **d-f,** Adipose tissue measurements on hematoxylin-eosin (H&E)-stained adult zebrafish sagittal sections. (**d**) Adipose tissue area per total section area (average of three sections/biological replicate) and (**e**) adipocyte size (average of 20–30 adipocytes of ventral adipose tissue per biological replicate) (Δ1 +/+ n=3, Δ2 n=8). SCAF1^Δ1^ and SCAF1^Δ2^ are represented with circles and squares, respectively. (**f**) Representative images of ventral fat deposits (dotted lines). **g**, Number of eggs per clutch (standard diet Δ1 +/+ n=8, Δ1 -/- n=8, Δ2 +/+ n=7, Δ2 -/- n=12, double diet Δ1 +/+ n=7, Δ1 -/- n=8, Δ2 +/+ n=8, Δ2 -/- n=9). **h**, Representative images of H&E-stained ovaries. Black dotted lines outline the ovaries, blue dotted line indicates a mature follicle. **i**, Quantification of mature ovary follicles per ovary section (average of three sections/biological sample) (Δ1 n=2-3, Δ2 n=6). **b, c**, Two-way ANOVA, **d, h**, unpaired t- test, **e,f**, one-way ANOVA. Data are represented as mean ± SD. * p<0.05, ** p<0.01, *** p<0.001, **** p<0.0001. Scale bars = 500 μm.

To understand the underlying adaptation of mutants and the positive effect of the double diet, we tested whether this phenomenon could be reproduced by feeding the animals with a diet with double the amount of fat (high-fat diet, HFD), which increases the caloric content to 141%. Under HFD, the differences in growth and fertility induced by the lack of SCAF1 remained, showing that it is the double caloric intake mostly provided by proteins rather than fats that is responsible for rescuing growth and weight in SCAF1^-/-^ animals (Fig 6).

**Figure 6.**
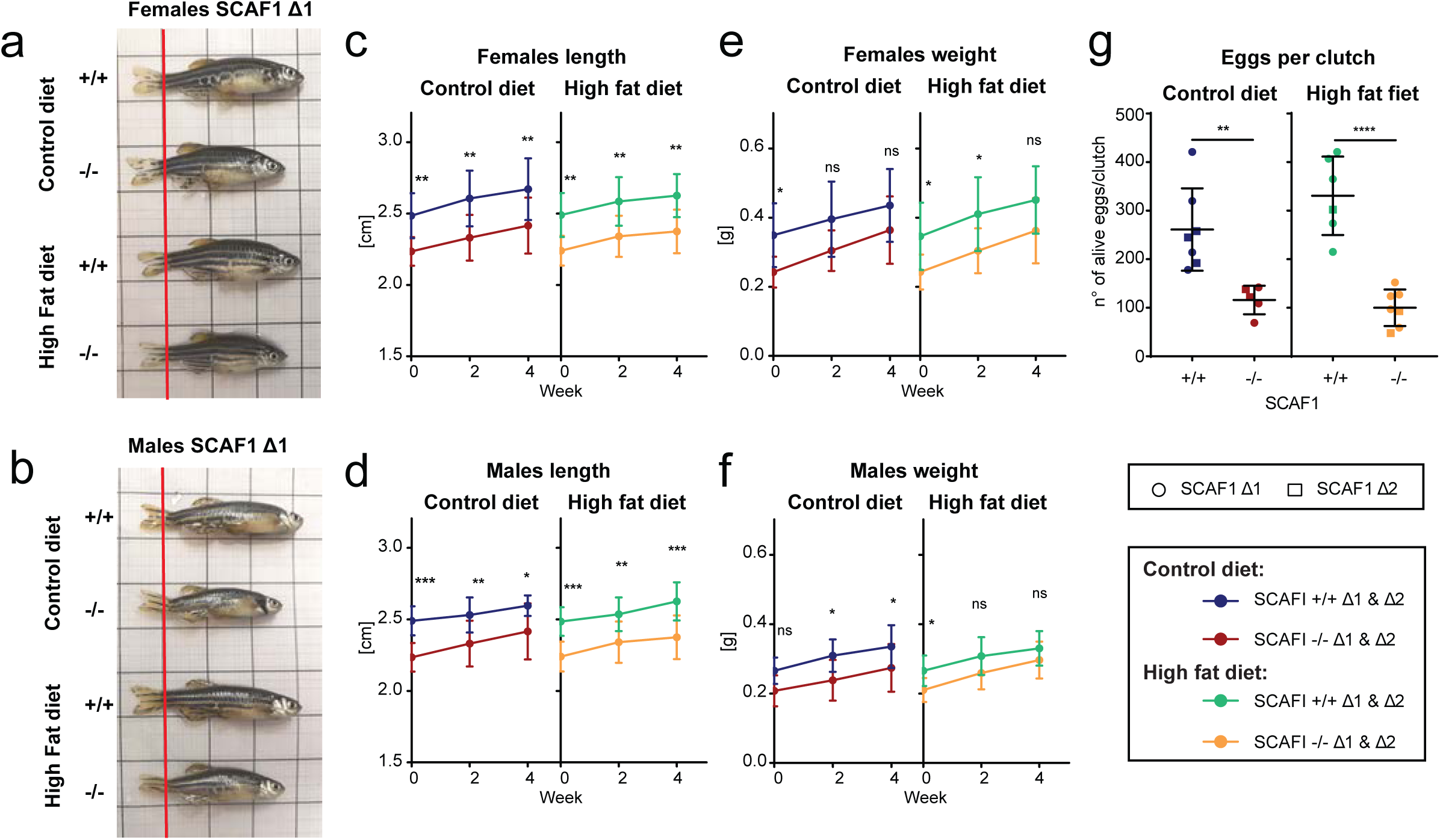
Lack of recovery of SCAF1^-/-^ phenotypes after high-fat diet. **a, b,** Representative images of SCAF1^-/-^ and SCAF1^+/+^ females (**a**) and males (**b**) fed with the indicated diets. **c, d,** Length of females (**c**) and males (d) after the indicated diets. **e, f,** Weight of females (**e**) and males (**f**) after the indicated diets (Δ1 +/+ n=5, Δ1 -/- n=5, Δ2 +/+ n=5, Δ2 -/- n=5). **g**, number of eggs per clutch (standard diet: Δ1 +/+ n=4, Δ1 -/- n=3, Δ2 +/+ n=3, Δ2 -/- n=2, double diet: Δ1 +/+ n=5, Δ1 -/-, n=5 Δ2 +/+ n=1, Δ2 -/- n=2). Two-way ANOVA. Data are represented as mean ± SD. * p<0.05, ** p<0.01, *** p<0.001, **** p<0.0001.

We next aimed to analyze whether diet could influence complex assembly or bioenergetics in the absence of SCAF1 (Fig. 7). Analysis of BNGE gels from double diet and standard diet wild-type and mutant fish revealed no differences in the super-assembly pattern of CIII and CIV (Fig. 7a). Interestingly, the double diet significantly increased the maximum respiration (site I+II) capacity of isolated adult mitochondria from SCAF1^-/-^ animals, but it remained below wild-type respirations levels (Fig. 7b). Therefore, despite the phenotype recovery, OXPHOS capacity in double diet is not fully normalized.

**Figure 7.**
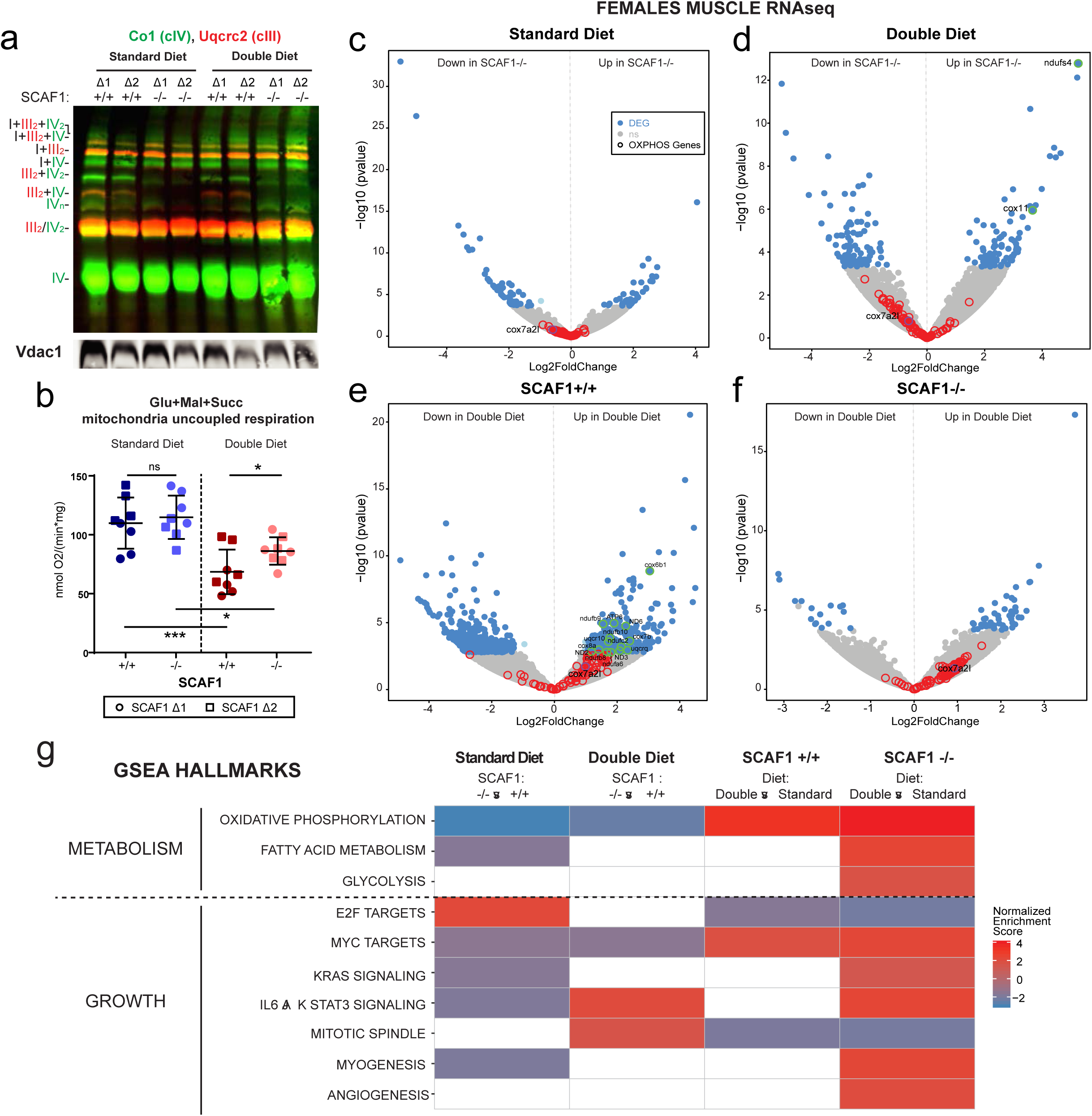
Molecular basis for diet-induced recovery of SCAF1^-/-^ phenotypes. **a,** Immunoblot of the indicated proteins of BNGE from female SCAF1^+/+^ and SCAF1^-/-^ whole zebrafish mitochondria for the indicated diet (representative of n=2). **b** Maximum uncoupled (FCCP) oxygen consumption rate in whole zebrafish mitochondria (females Δ1 n=4 and Δ2 n=4, and same number for their respective controls) with glutamate (Glu), malate (Mal) and succinate (Succ). **c-g** RNAseq data from SCAF1^+/+^ and SCAF1^-/-^ skeletal muscle for the indicated diet (standard diet SCAF1^+/+^ and SCAF1^-/-^ n=4, double diet SCAF1^+/+^ n=3, SCAF1^-/-^ n=4). **c-f** Volcano plots of differentially expressed genes (DEGs). (**c**) Comparison between SCAF1^-/-^ and SCAF1^+/+^ zebrafish in standard diet. (**d**) Comparison between SCAF1^-/-^ and SCAF1^+/+^ zebrafish in double diet. (**e**) Comparison of SCAF1^+/+^ zebrafish in double diet and standard diet. (**f**) Comparison of SCAF1^-/-^ zebrafish in double diet and standard diet. In blue, significant DEGs (p-adj < 0.05, log_2_FC > |1|), in grey not significant DEGs, red circles represent non-significant differentially regulated OXPHOS genes, green circles represent significant differentially regulated OXPHOS genes, purple SCAF1 (*cox7a2l*). (**g**) Heatmap of differentially regulated hallmarks in Gene Set Enrichment Analysis (GSEA) in the indicated comparisons. **b**, T-test analysis. Data are represented as mean ± SD. * p<0.05, ** p<0.01, *** p<0.001, **** p<0.0001.

To gain molecular insight into the mechanism of the body size recovery upon double diet, we performed RNAseq on female skeletal muscle under the different diet regimens. Most of the OXPHOS genes were slightly downregulated in SCAF1^-/-^ muscle compared with SCAF1^+/+^ muscle, both in standard diet (Fig. 7c) and in double diet (Fig. 7d) conditions. In double diet, skeletal muscle showed an increase in the expression of OXPHOS genes both for SCAF1^+/+^ (Fig. 7e) and SCAF1^-/-^ (Fig. 7f) animals, although this increase was not sufficient to recover OXPHOS gene levels to those of SCAF1^+/+^ animals (Fig. 7d). Consistent with these findings, the OXPHOS gene set enrichment analysis (GSEA) hallmark was downregulated in SCAF1^-/-^ muscle independently of the diet. By contrast, the fatty acid metabolism GSEA hallmark was downregulated in SCAF1^-/-^ muscle only in the standard diet condition (Fig. 7g). Double diet induced an upregulation of fatty acid metabolism and glycolysis GSEA hallmarks in SCAF1^-/-^ fish, which could be responsible for the recovery of the fish size. Indeed, analysis of SCAF1^-/-^ fish gene expression under standard diet revealed downregulation of pathways related to growth, such those involved in myogenesis, KRAS and IL6 JAK/STAT3 signaling pathways^30^. In agreement with this, the aforementioned pathways were those most strikingly upregulated in SCAF1^-/-^ muscle upon double diet. In addition, the hallmark E2F targets involved in G1/S transition in the cell cycle (cell cycle arrest) were recovered with the double diet in SCAF1^-/-^ muscle (Fig. 7g). Overall, the transcriptomic analysis suggests that rather than restoring OXPHOS super-assembly capacity, double diet leads to the activation of the affected OXPHOS maximum capacity accompanied by the enhancement of a variety metabolic pathways, allowing the rescue of growth in SCAF1^-/-^ zebrafish.

## DISCUSSION

Three major conclusions can be drawn from the results presented in this report: (1) the exhaustive description of the super-assembly pattern of the respiratory complexes in zebrafish corroborates the ***plasticity model*** of mitochondrial ETC organization^31^; (2) the role of SCAF1 as a SC assembly factor responsible for CIII-CIV interaction is conserved in non-mammalian vertebrates and; (3) the physiological and bioenergetic role of super-assembly in the optimization of metabolic resources and in particular of SCAF1 has been demonstrated.

Although the organization pattern of respiratory SC in zebrafish assessed by BNGE is very similar to that described in mammals, it reveals some unique features. The more remarkable of these is the higher proportion of super-assembly of the dimer CIV (IV_2_), both in the form of III_2_+IV_2_ and I+III_2_+IV_2_ SCs. This contrasts with the arrangement of the SC between III and IV in yeast, which is formed by the association of a dimer of CIII with two monomers of CIV instead of a dimer^32^. Also of interest is the presence of the SCs I+IV and I+IV_2_, for which there is only one report in mammals based on mass spectrophotometry^6^, and that is confirmed in zebrafish in the present study. The presence of this association is intriguing since the lack of CIII precludes a role in respiration. It might only be a consequence of the mitochondrial inner membrane solubilization process without any relevance *in vivo* or, as occurs with the multimerization of CV^33^, it might have a structural role. This is of importance to avoid the incorrect interpretation of BNGE data. Indeed, given that I+IV_2_ and I+III_2_ have similar mass and co-migrate, they may be wrongly identified as a respirasome. Finally, we describe here a truncated form of SCAF1, which we have found also in the mouse (manuscript in preparation), and that is generated by the proteolytic digestion by calpain 1 (manuscript in preparation). The physiological relevance of this phenomenon is still unclear. In any case, it is very relevant to correctly interpret the BNGE profiles. The fact that this precise processing is observed both in zebrafish and mouse SCAF1 calls for a deeper investigation of this phenomenon.

SCAF1^-/-^ zebrafish lack III_2_+IV, III_2_+IV_2_ and I+III_2_+IV_2_ SCs, but maintain a substantial amount of I+III_2_+IV. The presence of respirasomes persisting in the absence of functional SCAF1 in some mouse tissues^2, 34, 35^ and human cell lines^17, 18^ might be explained by their ability to interact independently with CI even in the absence of a physical interaction between them^7, 36^. The fact that we observed SCs containing I+IV or I+III_2_ by BNGE supports the idea of a respirasome assembled by interactions of CIII and CIV solely with CI in the SCAF1^-/-^ animals. Nevertheless, quantitative estimations revealed that the ablation of SCAF1 reduced the amount of respirasomes, mimicking our findings in mice^2^.

The potential physiological and bioenergetic role of respiratory SCs and SCAF1 remained an open question^17, 18^. Here, we show the impact of the ablation of SCAF1 on SC assembly and on the substrate-dependent electron flux. Thus, in the presence of SCAF1, neither CI-dependent respiration (site I) nor CII-dependent respiration (site II) reaches their maximum rates obtained by their combination (site I+II). By contrast, CII respiration was able to reach its maximum rate in the absence of SCAF1. This observation reproduced, at the organismal level, the impact of SCAF1 in isolated mitochondria from mouse liver^1^ and human cell lines^18^ (Fig. 8).

**Figure 8.**
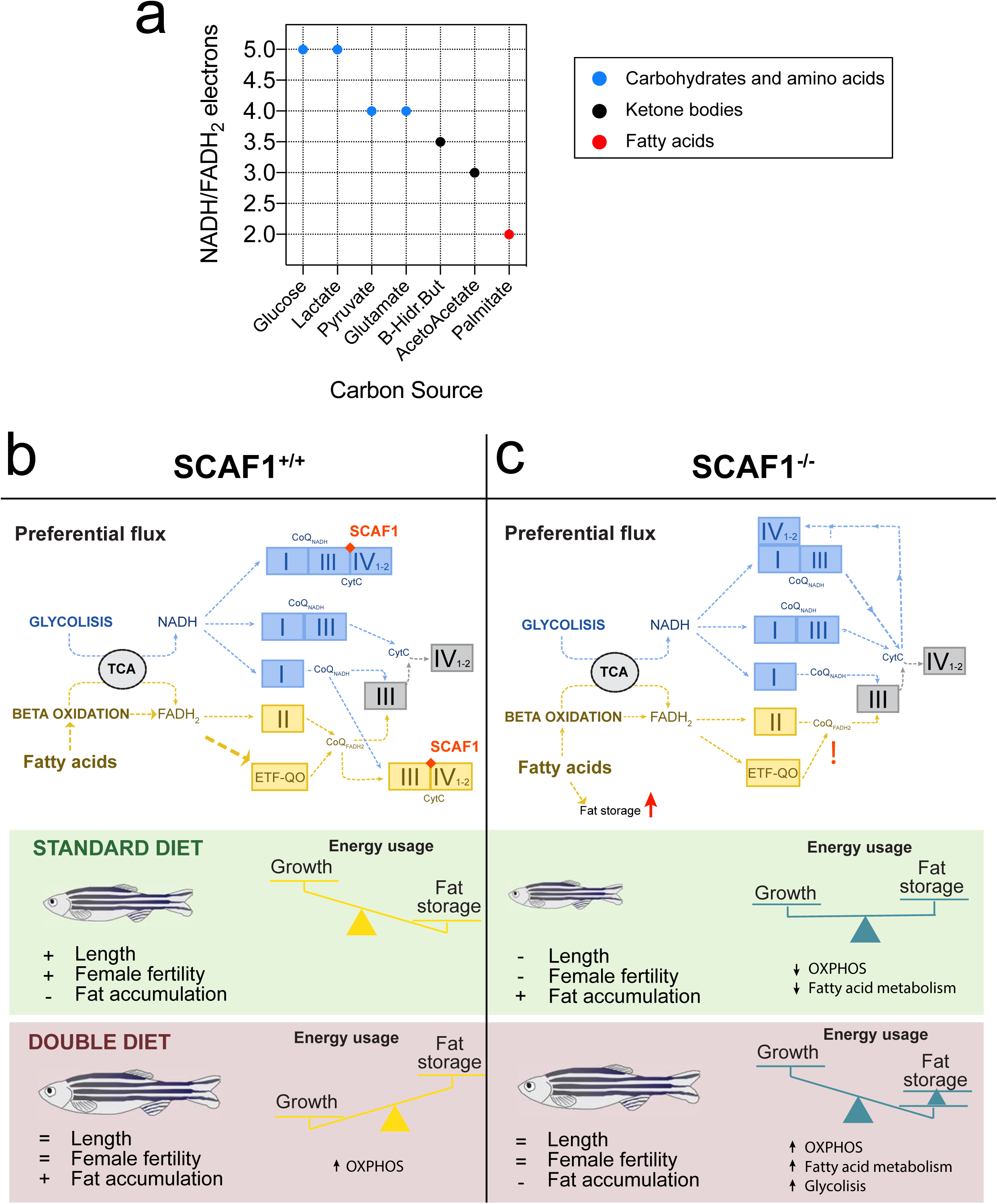
Schematic representation of molecular and physiological phenotypes of SCAF1^+/+^ and SCAF1^-/-^. **a,** Ratio of reduction equivalents in the form of NADH and FADH_2_ for different substrates. **b**,**c,** Model of the organization of the electron flux in wild-type (b) and SCAF1 deficient (c) fish showing the summary of the phenotypic consequences of standard or double diet. ETF-QO = electron transfer flavoprotein-ubiquinone oxidoreductase.

Our assessment of the physiological impact of SCAF1 ablation demonstrates its role and, by extension, the role of respiratory SCs, in the bioenergetic fine-tuning of mitochondria. The lack of SC formation due to SCAF1 ablation reduce the efficiency of conversion of nutrients into energy, mimicking some features associated with malnutrition in zebrafish, such as growth and female fertility, despite a normal food intake^23^. In agreement, overfeeding compensates the phenotypic effects of SCAF1^-/-^ animals. It might seem paradoxical that there is an increase in adipose tissue deposits in normal diet-fed SCAF1^-/-^ zebrafish compared with controls. Moreover, both groups showed increased adiposity under double diet, but more so in wild-type animals. All these features indicate that the flexibility of the ETC to adapt to different fuel sources is compromised in SCAF1^-/-^ zebrafish, particularly for the use of fatty acids. This also provides a plausible explanation for their inability to recover growth under HFD, and explains why a double diet, containing additional nutrients such as proteins, is needed for phenotypic rescue. To understand why the lack of super-assembly between complexes III and IV could be relevant for the proper adaptation to the fuel type, one has to consider that the catabolism of different carbon sources generates reducing equivalents in the form of NADH and FADH_2_ that are oxidized by the ETC. Different carbon sources generate a characteristic proportion between NADH/FADH_2_, ranging from 5 for glucose to 2 for fatty acids such as palmitate (Fig. 8a). NADH electrons enter the ETC through CI, which delivers them to CoQ. FADH_2_ electrons skip CI and are delivered directly to CoQ (Fig 8b). We previously proposed that the role of ETC super-assembly is to organize the flux of electrons to prevent competition between the two gateways for electrons by the segmentation of CoQ, CIII, cyt c and CIV in different subpopulations with preferential commitment either to NADH- or FADH_2_-derived electrons, and a third subpopulation that can be shared by both^1, 37^ (Fig 8b). We found in mice that the super-assembly is dynamic in response to different carbon sources and that this reorganization prevents the elevation of reduced CoQ and the increase in ROS^9^. The lack of SCAF1 prevents the segmentation of the CIV, eliminating the fraction preferentially dedicated to FADH_2_ electrons (Fig 8c). Since fatty acid oxidation is more demanding than pyruvate or glutamate oxidation (glucose or amino acids) for the FADH_2_ pathway, its oxidation will lose performance in the absence of SCAF1 (Fig 8c). Under normal diet the organism will sense normal levels of feeding but since SCAF1^-/-^ mitochondria cannot reach an optimized structure, neither sugar, amino acids nor lipids will be catabolized at an optimal rate. The fatty acid oxidation rate in SCAF1^-/-^ animals will be more affected, leading to elevated levels of circulating fat that will trigger their delivery to adipocytes. Under double diet the excess of nutrients triggers an overall increase in the catabolic capacity of both wild-type and mutant fish, as revealed by RNAseq, particularly in OXPHOS and lipid metabolism pathways, causing a modest elevation in OXPHOS capacity in SCAF1^-/-^ animals. This induces both enhanced degradation and storage of lipids. However, although the super-assembly between complex III and IV is not restored, the modest increase in OXPHOS capacity in combination with the activation of fatty acid metabolism allows normal growth to be achieved. Therefore, nutrition impacts the phenotypic manifestations of SCAF1^-/-^ zebrafish.

In summary, the incorporation of CIV into SCs through SCAF1 confers an adaptive advantage under limited nutrient availability and provides enhanced fitness to optimize OXPHOS performance.

## Acknowledgements

We are grateful to M. M. Muñoz-Hernandez, Raquel Martínez de Mena and Dr Concepción Jiménez for technical assistance; the Animal facility and Microscopy Units from CNIC and R. Baal for fish husbandry; and Sandra Nansoz from the Department of Intensive Care Medicine for use of the Oroboros equipment at the University of Bern and Inselspital. Electron microscopy sample preparation and imaging were performed with devices supported by the Microscopy Imaging Center of the University of Bern. J.A.E. was supported by Spanish Ministry of Economy and Competitiveness, MINECO (SAF2015-65633-R), CIBERFES (CB16/10/00282) and HFSP (RGP0016/2018). N.M. was funded by the ERC starting grant 337703, HFSP (RGP0016/2018) and SNF 31003A-159721. J.V. was funded by MINECO (BIO2015-67580-P), by Carlos III Institute of Health-Fondo de Investigación Sanitaria (PRB3, IPT17/0019 - ISCIII-SGEFI / ERDF, ProteoRed), the Fundación La Marato TV3 and by “La Caixa” Banking Foundation (HR17-00247). The CNIC is supported by the Ministry of Economy, Industry and Competitiveness (MEIC) and the Pro-CNIC Foundation, and is a Severo Ochoa Center of Excellence (MEIC award SEV-2015-0505).

## Methods

### Protein sequences alignment

FASTA files of amino acids sequences of SCAF1 were obtained from the public NCBI databases for mouse *Cox7a2l*^113^ and *Cox7a2l*^111^. The zebrafish paralog (named *cox7a2l* or *cox7a3*) was identified in Ensembl and its amino acid sequence was aligned to mouse sequences using the Clustal Omega platform.

### Generation of SCAF1 loss of function fish and genotyping

CRISPR/Cas9 sgRNAs (sgRNA 1: GATCATAGCAGGGGATTCGG AGG, sgRNA2: GGAGTACATGGGTAAAAACA GGG) were designed using the CCTop^38^ tool, cloned in the plasmid MLM3636 (Addgene #43860), synthesized and purified as described elsewhere^39^.

The two sgRNAs were co-injected at 120–150 ng/μl together with homemade 6.5 μM Cas9 protein-produced from the pCS2-nCas9n plasmid (Addgene #47929) as described^40^ in NEB Cas9 buffer (NEB #B0386A) into zebrafish 1-cell-stage embryos.

Founder animals were identified at three months post-fertilization (mpf) by fin clip PCR analysis using the primers 5’TCCACTCTGCTTACTTCACAC3’ and 5’TTTGCTTTGTCTGTATGTCCTG3’ and were crossed with AB wild-type fish to generate F1 progeny. PCR products from the mutant allele of the F1 heterozygous were purified from gel bands (NEB #T1020S) and analyzed by Sanger sequencing. Two lines derived from different injection rounds and progenitors with two different deletions were established: SCAF1^Δ1^ and SCAF1^Δ2^ (deposited in Zfin as *cox7a3^ube1^* and *cox7a3^ube2^*, respectively).

Genotyping during line maintenance used the described primers. All experiments were performed comparing SCAF1^Δ1/Δ1^ and SCAF1^Δ2/Δ2^ with their respective sibling lines *cox7a3^+/+^* coming from the same founder and AB mating. A maximum of 4 in-cross generations were used for the experiments.

### Zebrafish husbandry

Experiments were approved by the Community of Madrid “*Dirección General de Medio Ambiente*” in Spain and the “*Amt für Landwirtschaft und Natur*” from the Canton of Bern, Switzerland. All animal procedures conformed to EU Directive 86/609/EEC and Recommendation 2007/526/EC regarding the protection of animals used for experimental and other scientific purposes, enforced in Spanish law under Real Decreto 1201/2005. Experiments in Switzerland were conducted under the licenses BE95/15 and BE11/17. Experiments were conducted with adult zebrafish aged 5–10 months and raised at 13 fish per 2 l tank. Housing conditions were 28°C temperature, 650–700 µs/cm conductivity, pH 7.5; lighting conditions were 14:10 hours (light: dark) and 10% water exchange per day. To guarantee a stable fish density between groups, occasionally SCAF1^-/-^, SCAF1^+/-^ or SCAF1^+/+^ fish were grown in the same tank as transparent Casper fish^41^.

### Mouse husbandry

As with zebrafish, all animal procedures conformed to EU Directive 86/609/EEC and Recommendation 2007/526/EC regarding the protection of animals used for experimental and other scientific purposes, enforced in Spanish law under *Real Decreto 1201/2005.* Four-month-old mice from the strains C57BL/6JOlaHsd and CD1 were used as a source for mitochondria purification of SCAF1^111^ and SCAF1^113^ mice, respectively. Original parental mice were purchased from Harlan Laboratories.

### Mitochondria isolation

Whole fish or lateral skeletal muscle were cut into small pieces and rinsed using PBS and homogenization medium A (0.32 M sucrose, 1 mM EDTA, and 10 mM Tris-HCl, pH 7.4) at 4°C. Clean minced tissue was transferred to a manual dounce tissue grinder containing homogenization medium A supplemented with fatty acid-free 0.1% bovine serum albumin (BSA-FFA) (Sigma A7030). Occasionally, a motor-driven Teflon pestle with 6 up and down strokes at 650 rpm was used to replace the manual tissue grinder. The homogenate was centrifuged twice for 10 min at 800 × *g* and 4°C. The supernatant was then centrifuged for 10 min at 10000 × *g* and 4°C. The pellet from this step was resuspended in 1 ml of MAITE buffer (0.25 mM sucrose, 75 mM sorbitol, 100 mM KCl, 0.05 mM EDTA, 5 mM MgCl_2_, 10 mM Tris-HCl and 10 mM ortophosphoric acid, pH.7.4) supplemented with 0.1% BSA-FFA and centrifuged for 10 min at 10000 × *g* at 4°C. The clean pellet was resuspended in 0.5–1 ml MAITE+BSA-FFA buffer and the concentration was quantified using the Bradford assay (Sigma-Aldrich). Mitochondria were stored at -80°C in MAITE+BSA-FFA buffer or centrifuged for 10 min at 10000 × *g* and 4°C and subsequently resuspended in the buffer required for the following analysis. All steps were performed on ice.

For functional mitochondria assays, clean minced tissue was homogenized in homogenization buffer (67 mM sucrose, 50 mM Tris, 50 mM KCl, 10 mM EDTA, 0.1% BSA-FFA, pH 7.4)^42^ and centrifuged twice for 10 min at 800 × *g* and 4 °C. The supernatant was centrifuged 10 min at 8000 × *g* and 4 °C and washed with MAITE buffer as described above.

Mouse mitochondria were isolated from soleus skeletal muscle or liver as described^43^, solubilized in 4 g/g digitonin and run in parallel with zebrafish samples.

### SDS and blue native gel electrophoresis

A total of 100–200 µg of whole fish mitochondria were resuspended in loading buffer (50 mM Tris-HCl pH 6.8, 2% SDS, 10% glycerol, 1% β-mercaptoethanol, 12.5 mM EDTA, 0.02% bromophenol blue) at a final concentration of 2 µg/µl, and 30 µg were loaded onto 12% hand-cast sodium dodecyl sulfate (SDS) acrylamide gels.

BNGE was performed as described^44^ using 4 g/g digitonin-treated mitochondria; 30 µg of muscle or whole fish mitochondria or 20 µg of mouse soleus muscle or liver mitochondria were loaded onto 15 well 3–13 % hand-cast native gels.

For 2D-BNGE, the first dimension used 125 µg whole-body zebrafish mitochondria treated with 4 g/g digitonin in 10-well 3–13% hand-cast native gels. The second dimension was performed in native 3–13% gels adding 0.01% DDM in the electrophoresis cathode buffer^44^.

Two-dimensional denaturing electrophoresis (2D-BNGE/SDS-PAGE) was performed using 125 µg whole body zebrafish mitochondria treated with 4 g/g digitonin in 10-well 3–13 % hand-cast native gels as the first dimension. The second denaturing dimension with SDS was performed as described^44^.

### BNGE in-gel activity assays

After electrophoresis, BNGE gels were incubated for CI (0.1 mg/ml NADH and 2.5 mg/ml nitroblue-tetrazolium in 2 mM Tris-HCl pH 7.4 buffer) or CIV (1 mg/ml cytochrome C and 0.5 mg/ml in 3,3’-diaminobenzidine in phosphate buffer pH 7.4) activity^45^.

### Immunoblotting

BNGE or SDS gels were electroblotted onto Hybond-P-polyvinylene fluoride (PVDF) membranes (Immobilon-FL, IPFL00010) and immunoblotted with antibodies against the different subunits of the OXPHOS complexes: anti-Ndufs3 mouse monoclonal (Abcam, AB14711), anti-Uqcrc2 rabbit polyclonal (Proteintech, 14742-1-AC), anti-Co1 mouse monoclonal (ThermoFisher, 459600) and anti-Cox7a2l rabbit polyclonal (St John’s laboratory, STJ110597). Anti-Vdac1 (Abcam, AB15895) was used as loading control.

Secondary antibodies used were anti-rabbit IgG (H+L) Alexa Fluor 680 conjugate (Life Technologies, A-21076) and anti-mouse dylight 800 (Rockland, 610145002) and images were acquired with the ODYSSEY Infrared Imaging System (LI-COR). Immunoblotting of 1-dimension SDS using COX7A2L antibody were analyzed by the ECL detection system using polyclonal goat anti-rabbit (Dako, P0448) as the secondary antibody.

### Low Proteins/Low Fats (LP/LF) diet experiment

One experiment was performed with 8 wild-type 7 mpf fish per group at a density 5 fish/l (transparent Casper fish^41^ were used to reach tank density). Fish were randomly selected, measured at the beginning of the experiment and assigned to one diet group. The control group was fed with Sparos Control diet and the experimental group was fed with Sparos LP/LF diet (Supplementary data Table 1) three times/day during week days and one time/day during weekends (15 mg/tank per dose). Fish were measured and sacrificed after 6 weeks. Mitochondria were freshly isolated in pools of 2 animals/sample.

### Blue-DiS proteomics

Three replicates of 200 µg of 4 g/g digitonin-treated whole fish mitochondria from SCAF1^-/-^ and SCAF1^+/+^ animals were run in a 3–13 % hand-casted native gels, which were then stained with Coomassie Brilliant Blue R-250 and cut in 36 slices. The slices were processed and analyzed by data-independent mass spectrometry as previously described^2^.

### Fish size, length and embryo laying assessment

Anesthetized fish were measured in length with a millimetric ruler and weighed on a precision balance. For fertility assessment, one female was crossed with one male of the same genetic background and eggs were collected 20 min after direct mating. The number of live eggs was manually counted after laying.

### Zebrafish swimming performance

Critical swimming speed (maximum swimming speed) was assessed in 7–9 mpf untrained fish in a swimming performance tunnel (Loligo Systems, #SY28000) by increasing the current by 5 cm/s every 2 min with an initial speed of 5 cm/s until the fish no longer kept their position and fatigued^46^. Critical swimming speed was assessed in a blinded manner.

### Zebrafish histology

Whole fish where fixed in 4% para-formaldehyde for 24 hours, washed three times in PBS for 10 min and decalcified in Immunocal (American MasterTech) at room temperature (RT) for 24 hours. Tissues where dehydrated and embedded in paraffin blocks. Histological sections (7 µm) were used for hematoxylin-eosin staining. Three sagittal sections of representative areas per biological replicate were analyzed in ImageJ/Fiji and the average was represented for each biological sample. Adipose tissue area was measured in the dorsal, ventral, visceral and intramuscular areas, summarized and divided by the total tissue area. The area of 20–30 adipocytes in the ventral fat were measured per biological sample.

### Transmission electron microscopy

Hearts from 5 mpf zebrafish were fixed with 2.5% glutaraldehyde (Agar Scientific, Stansted, Essex, UK) and 2% paraformaldehyde (Merck, Darmstadt, Germany) in 0.1 M Na-cacodylate-buffer (Merck), pH of 7.33. Samples were fixed for 24 hours before further processing. They were then washed with 0.1 M Na-cacodylate-buffer three times for 5 min, post-fixed with 1% OsO4 (Electron Microscopy Sciences, Hatfield, USA) in 0.1 M Na-cacodylate-buffer at 4°C for 2 hours, and then washed in 0.05 M maleic acid (Merck, Darmstadt, Germany) three times for 5 min. Thereafter, samples were dehydrated in 70, 80, and 96% ethanol (Alcosuisse, Switzerland) for 15 min at RT. Subsequently, cells were immersed in 100% ethanol (Merck) three times for 10 min, in acetone (Merck) twice for 10 min, and finally in acetone-Epon (1:1) overnight at RT. Next, samples were embedded in Epon (Sigma-Aldrich, Buchs, Switzerland) and left to harden at 60°C for 5 days. Sections were produced with an ultramicrotome UC6 (Leica Microsystems, Vienna, Austria): semithin sections (1 µm) were used for light microscopy and stained with a solution of 0.5 % toluidine blue O (Merck); ultrathin sections (75 nm) were used for electron microscopy. The sections, mounted on 200-mesh copper grids, were stained with uranyl acetate (Electron Microscopy Sciences) and lead citrate (Leica Microsystems) with an ultrastainer (Leica Microsystems). Images were taken in a blinded manner using an FEI Tecnai Spirit Electron Microscope and analyzed on ImageJ/Fiji, also in a blinded manner.

### Mitochondrial DNA content

Genomic DNA was extracted from muscle and heart of 8 mpf fish with the DNeasy Blood & Tissue Kit (Quiagen). Three nanograms of DNA was used for qPCR using primers (nDNA: 5’ATGGGCTGGGCGATAAAATTGG3’, 5’ACATGTGCATGTCGCTCCCAAA3’; mtDNA: 5’CAAACACAAGCCTCGCCTGTTTAC3’, 5’CACTGACTTGATGGGGGAGACAGT3’) and method described before^53^. Data are represented with the formula 2*2^(nDNA CT – mtDNA CT).

### Embryo OXPHOS toxicity

Embryos (4 dpf) from in-crosses of SCAF1^-/-^ (SCAF1^Δ1/Δ1^ and SCAF1^Δ2/Δ2^) fish and their respective SCAF1^+/+^ counterparts were treated for 4 hours at 28°C with 3–4 concentrations of OXPHOS inhibitors: CI, rotenone (Sigma, R8875); CII, 3-nitroproionic acid (Sigma, N5636); CIII, antimycin A (Sigma, A8674); H^+^-ATPase oligomycin (Sigma, O4876); coupling inhibitor, carbonyl cyanide 4-trifluoromethoxyphenylhydrazone (FCCP) (Sigma, C2920), in 0.05% ethanol E3 medium; and CIV, sodium azide (Sigma S2002) in E3 medium. Ten embryos were placed in a 12-well plate and treated in a final volume of 2 ml^26^. Nine biological replicates in three technical replicates per fish group were analyzed and represented as a proportion of survival. Cardiac arrest was used as lethality parameter.

### Spectrophotometric measurements of OXPHOS enzymes

Whole adult fish isolated mitochondria after a freeze-drawn cycle were used to measure the enzymatic activity of respiratory complexes as described: CI, CII, CIV, citrate synthase^47^ and CIII ^49^. Values of CI, CII, CIII and CIV were normalized by citrate synthase activity.

### Spectrophotometric measurement of citrate synthase for mitochondrial content estimation

Citrate synthase activity was measured from hearts and muscles homogenized in homogenization medium A at 4°C. Experiments were carried out in a Tris-based buffer (pH 8) containing 0.1% Triton X-100, 5,5’ -dithio-bis (2-nitrobenzoic acid) (DNTB) and Acetyl-Coa. Absorbance increase at 412 nm due to DNTB reduction was tracked for 4 min at 30°C after addition of oxalacetate. Data were normalized by protein content of the homogenate measured by Bradford assay (Sigma Aldrich).

### Whole embryo oxygen consumption rate

Mitochondrial function was determined with an XFe24 extracellular flux analyzer (Seahorse Bioscience). OCR was measured in dechorionated 48 hpf embryos. Embryos were staged (same size was corroborated) and placed one per well on an islet capture microplate filled with E3 egg water. The plate was incubated in an incubator without CO_2_ at 28°C for 30 min. After measuring baseline OCR as an indication of basal respiration, OCR was measured after an injection of 2 µM of FCCP to determine maximal respiration. Finally, 0.5 µM of antimycin A and 0.5 µM of rotenone was added to block mitochondrial respiration.

### Oxygen consumption in isolated mitochondria from adult fish

Fresh isolated mitochondria (100 µg/ml) were analyzed using a Clark-type electrode (Oxygraph O2k; Oroboros Instruments, Innsbruck, Austria) at 28°C in MirO5 medium^48^. CI-dependent oxygen consumption (site I) was measured with 10 mM pyruvate or 10 mM glutamate and 5 mM malate. CII-dependent oxygen consumption (site II) was measured with 10 mM succinate and 0.2 µM rotenone. CI- and CII-dependent oxygen consumption (site I+II) was measured with the same concentrations of pyruvate or glutamate, malate and succinate. The uncoupled state was reached with the inhibition of CV (H+-ATPase) with 4 ng/ml oligomycin and a titration of FCCP in 0.1–0.5 µM intervals until reaching the stable maximum OCR. Respiration was stopped with 30 mM sodium cyanide. The RCR (State3/State4) and P/O ratio were measured in a Clark-type electrode (Hansatech) with the aforementioned substrates plus 0.5 mM ADP. State 3 was calculated at the maximum OCR after ADP addition, and state 4 was calculated when ADP was consumed. The P/O ratio was calculated from the same measurement.

### Double diet experiments

Three independent diet experiments were performed with 7–9 mpf fish Control siblings and SCAF1^-/-^ fish in each experiment were born the same day and were grown in the same conditions. Experiment one (starting at 8.5 mpf) and two (starting at 7 mpf) were performed with SCAF1^Δ2^ and their respective SCAF1^+/+^ fish (n=10 fish per experimental group, per experiment). Experiment three (starting at 7 mpf) was performed with SCAF1^Δ1^ and their respective SCAF1^+/+^ fish (n=20 fish per experimental group). Fish were randomly selected and length and weight were measured. They were then distributed in equal groups according to their measurements. Ten fish from mixed sex where placed per tank (5 fish/l in 2 l tanks) and assigned to a diet category. Standard diet was established as the diet followed in the fish facility (weekdays: 2 times/day dry food 15 mg/tank Gemma Micro 300, 1 time/day artemia, weekends: 1 time/day dry food 15 mg/tank Gemma Micro 300) and double diet (weekdays: 4 times/day dry food 15 mg/tank Gemma Micro 300, 1 time/day artemia, weekends: 1 time/day dry food 15 mg/tank and 1 time/day 30 mg Gemma Micro 300). Fish were maintained under these conditions for 6 weeks, measuring length and weight every 2 weeks during the 4 first weeks of the experiment. At week 5, fish were in-crossed and the number of eggs laid per female was counted. Data from the three different experiments are represented together. Fish were sacrificed after 6 weeks. Fish from experiment two were used for histological analysis and BNGE. Fish from experiment 3 were used for histological analysis, BNGE and RNAseq.

### High-fat diet experiments

Two independent diet experiments were performed with 7 mpf fish, one with SCAF1 ^Δ1^ and the other with SCAF1^Δ2^ (n=10 per experimental group, 5 females, 5 males). Fish were randomly selected, length and weight were measured, and they were distributed in equal groups according to their measurements. Ten fish from mixed sex where placed per tank (5 fish/l in 2 l tanks) and assigned to a diet category. The control group was fed with Sparos Control Diet (14.1% fats). The high-fat diet group was fed with a customized diet by Sparos modified from their Control Diet containing double the amount of fats (30.1%) (Supplementary data Table 1). Both groups were fed 3 times/day on weekdays and 1 time/day on weekends. Fish were maintained in these conditions for 6 weeks measuring length and weight every two weeks during the 4 first weeks of the experiment. At week 5, fish were in-crossed and the amount of eggs laid per female was counted. Data from SCAF1^Δ1^ and SCAF1^Δ2^ were plotted and analyzed together. Fish were sacrificed after 6 weeks.

### RNA isolation and sequencing

Muscle from females SCAF1^Δ1^ after 6 weeks on the diets (standard diet SCAF1^+/+^ and SCAF1^-/-^ n=4, double diet SCAF1^+/+^ n=3, SCAF1^-/-^ n=4) were dissected and stored at -80°C. RNA was isolated using Trizol and purified (Zymo RNA Clean & Concentrator kit). RNA purity was evaluated using the Agilent Fragment Analyzer and used for the bar-coded library generation (RIN values: standard diet SCAF1^+/+^: 9.5 / 9.4 / 8.8 / 8.6, SCAF1^-/-^:9.5 / 9.2 / 9.4 / 7.4; double diet SCAF1^+/+^: 8.8 / 8.1 / 7.9, SCAF1^-/-^:9.1 / 9.3 /9.2 /9.1). Libraries were sequenced on the Illumina HiSeq 2500 System. RNA-seq data and related information have been deposited in the Gene Expression Omnibus database with accession nos. GSE133487.

### RNA-Seq bioinformatic analysis

Sequencing reads were pseudo-aligned to *Danio rerio* cDNA database (Ensembl build 11, release 94) using Kallisto^49^ version 0.45.0. Quality control was assessed using FastQC. Downstream analysis was performed in RStudio and differential expression between design groups was tested using DESeq2^50^, with log2 fold change shrinkage. Volcano plots and heatmaps were generated using the base graphics and ggplot2 package.

*Danio rerio* Gene Stable IDs were translated to *Mus musculus* Gene Stable IDs using biomaRt^51^ package and the overall gene expression was analyzed with Gene Set Enrichment Analysis (GSEA)^52^ using the Hallmarks collection as biological insight. Only data with adjusted p-value and FDR < 0.05 were considered significant and represented.

### Statistical analysis

Normal distribution was tested using D’Agostino-Pearson omnibus and Shapiro-Wilk normality tests. Outliers were identified using the ROUT test (Q = 1%) and when identified, they were not used in the statistical test. A t-test was used when comparing two groups and one- or two-way analysis of variance (ANOVA) was used when more than two groups were analyzed. ANOVA multiple comparison were performed by Sidak’s multiple comparison test when not indicated or Fisher’s LSD test when indicated. All data representations and statistical analyses were performed using GraphPad Prism 7.

---

**Figure S1.**
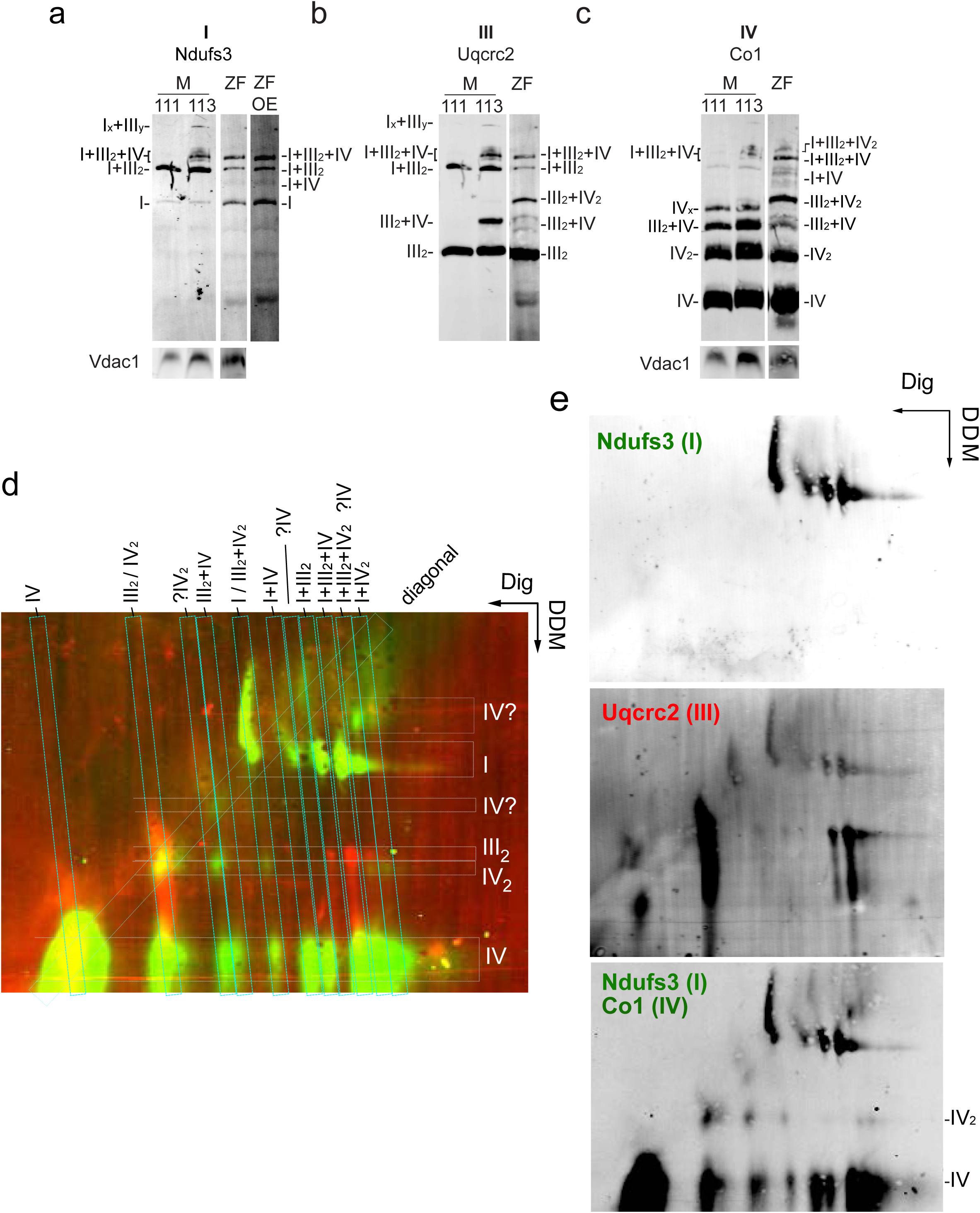
Characterization of OXPHOS super-assembly in zebrafish. **a-c,** Split channels of BNGE of mouse (M) C57BL/6J (111), CD1(113) and zebrafish (ZF) muscle mitochondria shown in Fig 1 a-b. **d,e**, Immunodetection of the indicated proteins after 2D: BNGE/DDM electrophoresis of whole-body zebrafish mitochondria (representative of n=3). Merged (**d**) and (**e**) split channels.

**Figure S2.**
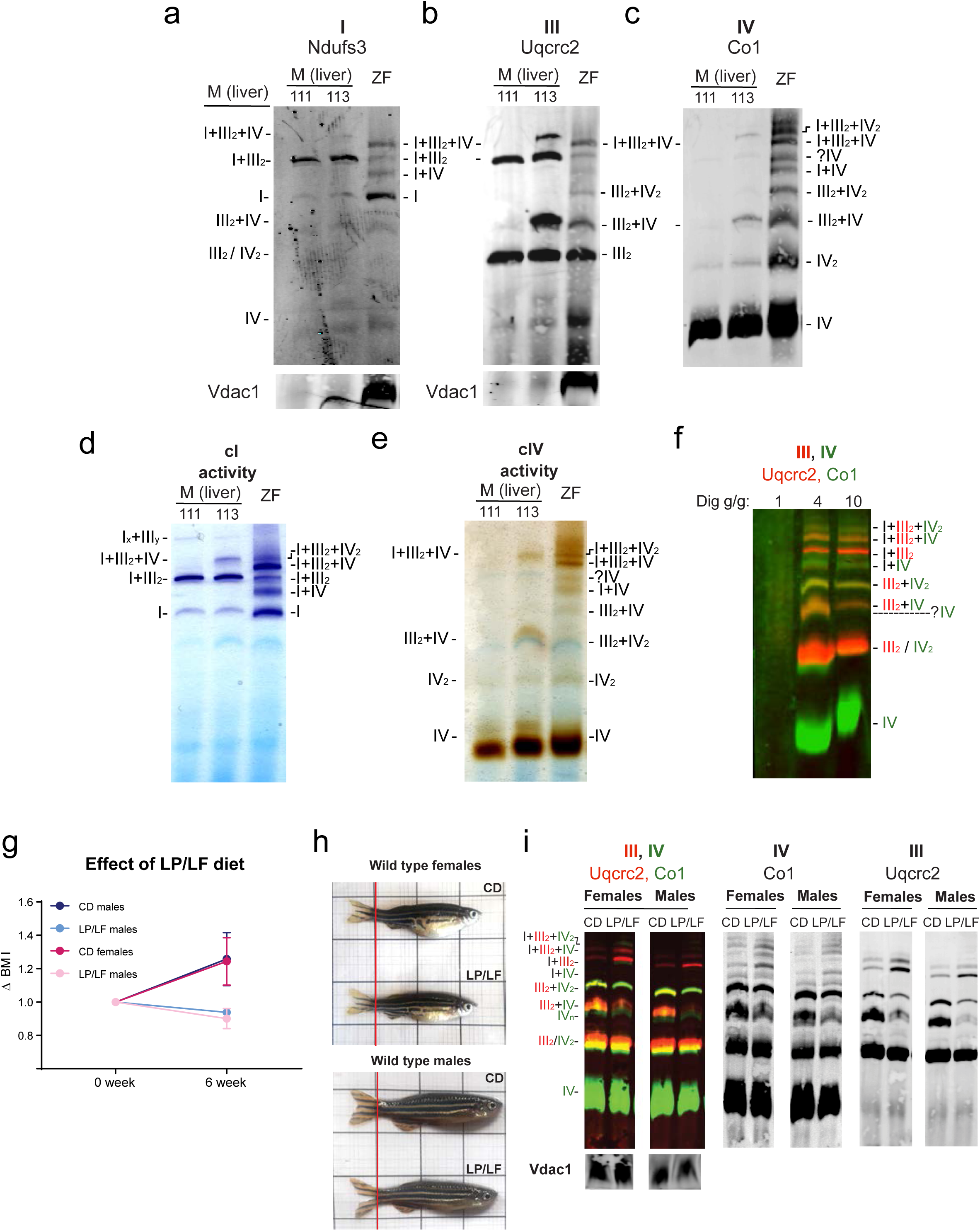
OXPHOS super-assembly in whole zebrafish and mouse liver in homeostasis and low protein/low fat diet. **a-e,** BNGE of mouse (M) C57BL/6J (111) and CD1(113) liver mitochondria and zebrafish (ZF) whole body mitochondria, digitonin-solubilized. (**a-c**) Immunodetection of the indicated proteins, (**d**) CI and (**e**) CIV in-gel activity (representative of two technical and three biological replicates). **f**, Immunodetection of the indicated proteins of digitonin-solubilized whole-body zebrafish mitochondria with different concentrations of digitonin. **g-i** BNGE of zebrafish fed with Low Proteins/Low Fats diet (LP/LF) and control diet. (**g**) variation of BMI of fish after 6 weeks in Low Proteins/Low Fats diet (LP/LF) and control diet (CD). (**h**) Representative images. (**i**) Representative BNGE of whole fish mitochondria of fish fed during 6 weeks in the indicated diet (experimental replicates n=2 are composed by a pool of n=2 biological replicates).

**Figure S3.**
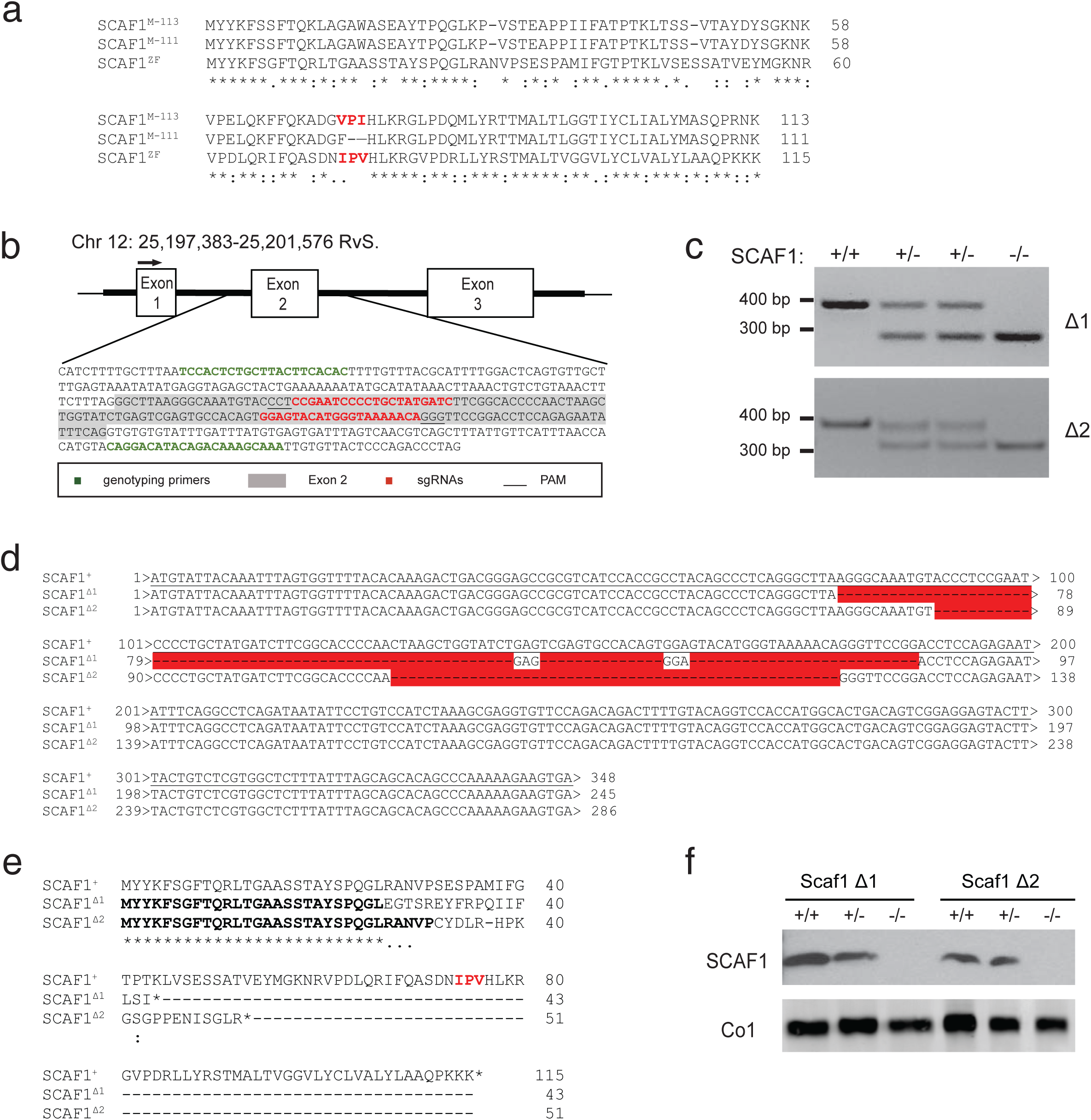
Generation of SCAF1 loss-of-function zebrafish models. **a**, Alignment of the amino acid sequences of mouse SCAF1^113^, SCAF1^111^ and zebrafish SCAF1 (Cox7a2l, Cox7a3). **b**, CRISPR/Cas9 target design for generation of zebrafish SCAF1 loss-of function-models. In green, primer sequences for genotyping, in red sgRNAs with their PAM sequence underlined. Exon sequences are highlighted in grey. **c**, Representative image of PCR products of the two established SCAF1 loss-of-function lines (SCAF1^Δ1^ and SCAF1^Δ2^). **d**, Sequence of SCAF1^Δ1^ and SCAF1^Δ2^ mutant alleles. In red, the deleted sequence. **e**, Predicted amino acid sequence of SCAF1^Δ1^ and SCAF1^Δ2^. **f**, Western blot of isolated mitochondria from SCAF1^Δ1^ and SCAF1^Δ2^ fish (representative of 4 biological replicates).

**Figure S4.**
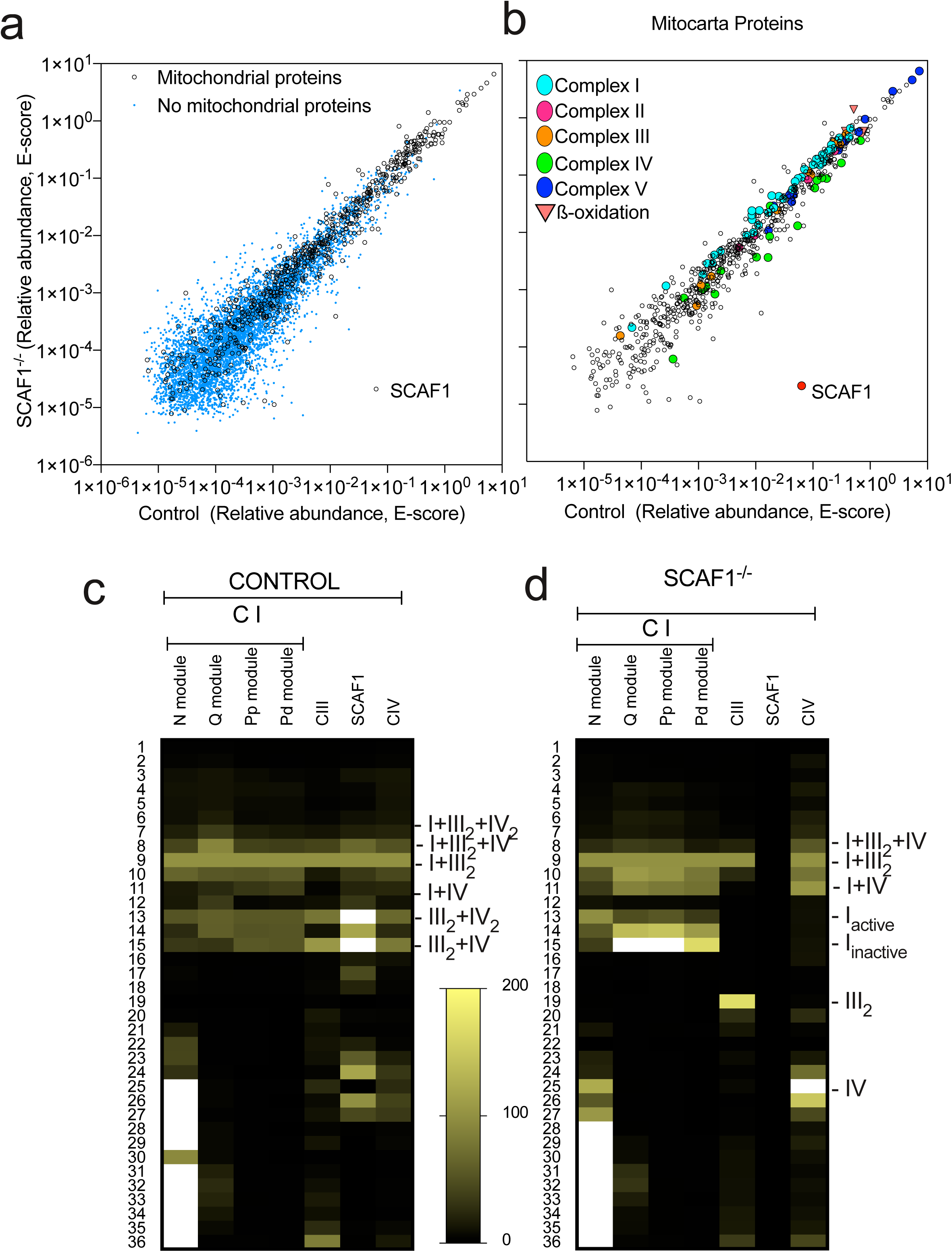
Analysis of mitochondrial complexes by Blue-DiS proteomics. **a** and **b**, Correlation between the abundance of proteins (expressed as sum of E-scores of the corresponding peptides) detected in the analysis of SCAF1^+/+^ and SCAF1^-/-^ animals. Proteins were considered mitochondrial according to the classification in the mouse Mitocarta 2.0 database. Non-mitochondrial proteins include true non-mitochondrial proteins and potential mitochondrial proteins that failed to be identified as such. In **b,** only the mitochondrial proteins are represented, indicating the proteins from the indicated groups. **c,d**, Heatmaps showing the summed absolute abundance of selected protein groups across BNGE gel slices. For a better comparison, absolute abundances were normalized using the values of slice 9 as a reference. Qualitative migration of the added E-score value for each indicated complex, subcomplex or protein. For each line, data were normalized within a 100 to 0 range, with 100 being the value of slice 9. The color scale is established as a linear increase from black (being 0) to the green in slice 9 (being 100). Any value over 100 is white.

**Figure S5.**
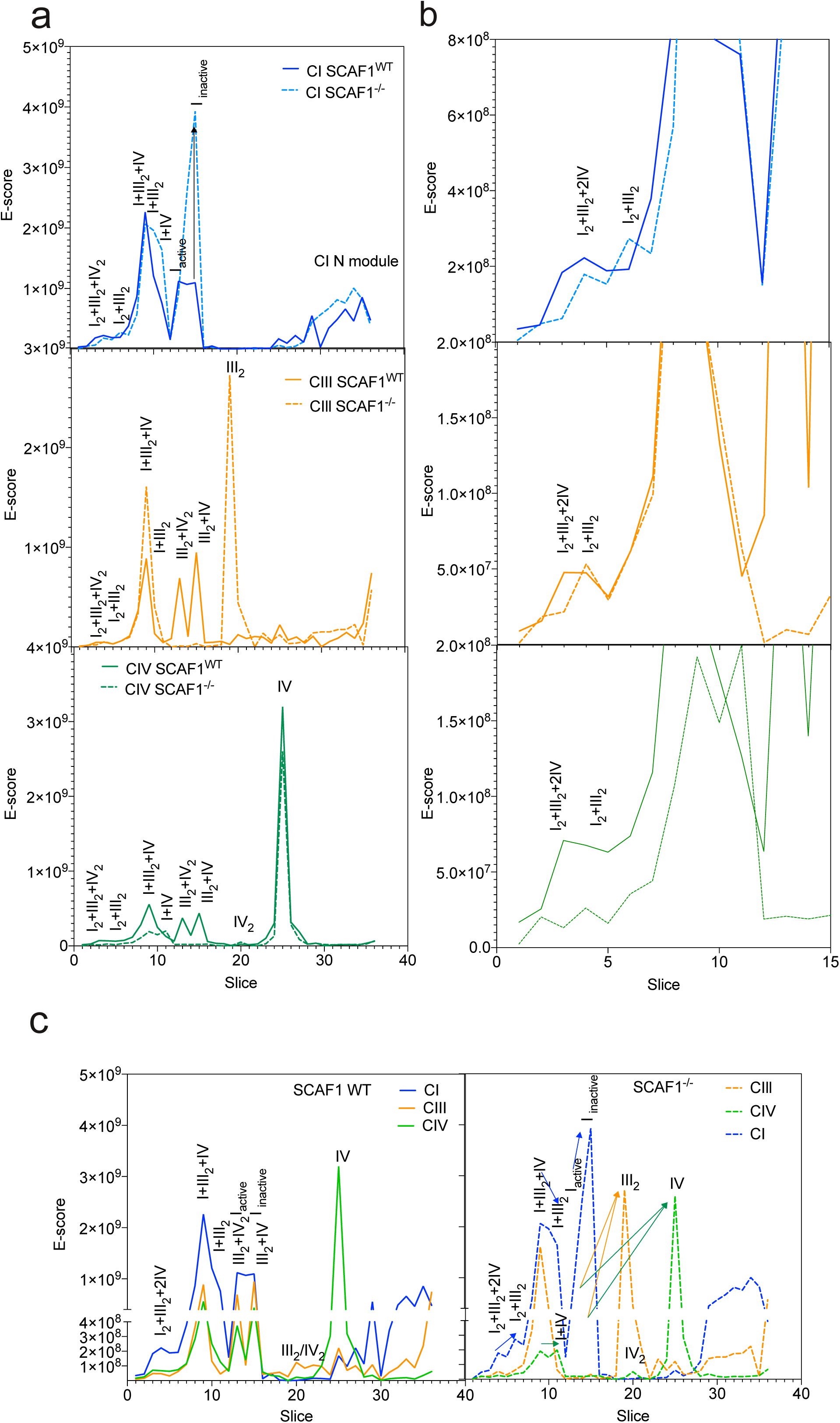
Analysis of the quantitative differences between Blue-DiS profiles of SCAF1^+/+^ and SCAF1^-/-^ animals. **a**, Differences in quantity profiles of CI-, CIII- and CIV-related complexes and SCs in control and SCAF1^-/-^ samples. The insets focus on the differences at very high molecular weights (slices 1 to 15). **b,** Comparative analysis of quantitative profiles of complexes and SCs in control or SCAF1^-/-^ samples. Arrows indicate increase, decrease or shifts of complexes observed between SCAF1^-/-^ and control samples.

**Figure S6.**
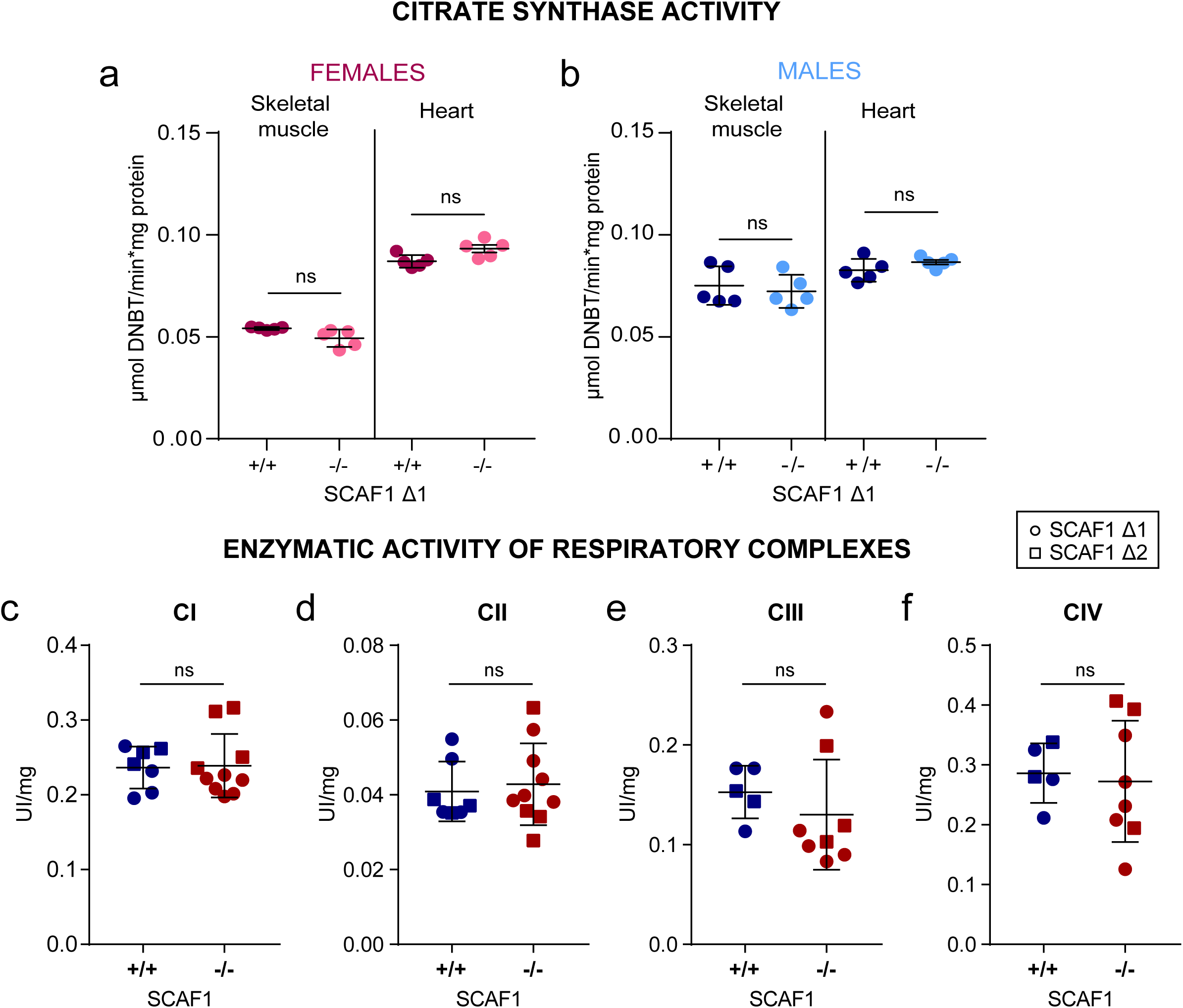
Spectrophotometrically measurements of mitochondrial content by citrate synthase activity and OXPHOS enzymes. **a,b** Citrate synthase activity of heart and muscle tissue homogenates in females (**a**) and males (**b**) adult zebrafish (Δ1 n=5 for each group). **c-f** Enzymatic activity of respiratory complexes in isolated mitochondria of whole adult zebrafish. (**c**) Complex I (cI) and (**d**) Complex II (cII) (Δ1 +/+ n=4, Δ1 -/- n=6, Δ2 +/+ n=3, Δ2 -/- n=4). (**e**) complex III (cIII) and (**f**) complex IV (cIV) (Δ1 +/+ n=3, Δ1 -/- n=5, Δ2 +/+ n=2, Δ2 -/- n=3). T-test analysis. Data are represented as mean ± SD. * p<0.05, ** p<0.01, *** p<0.001, **** p<0.0001.

**Figure S7.**
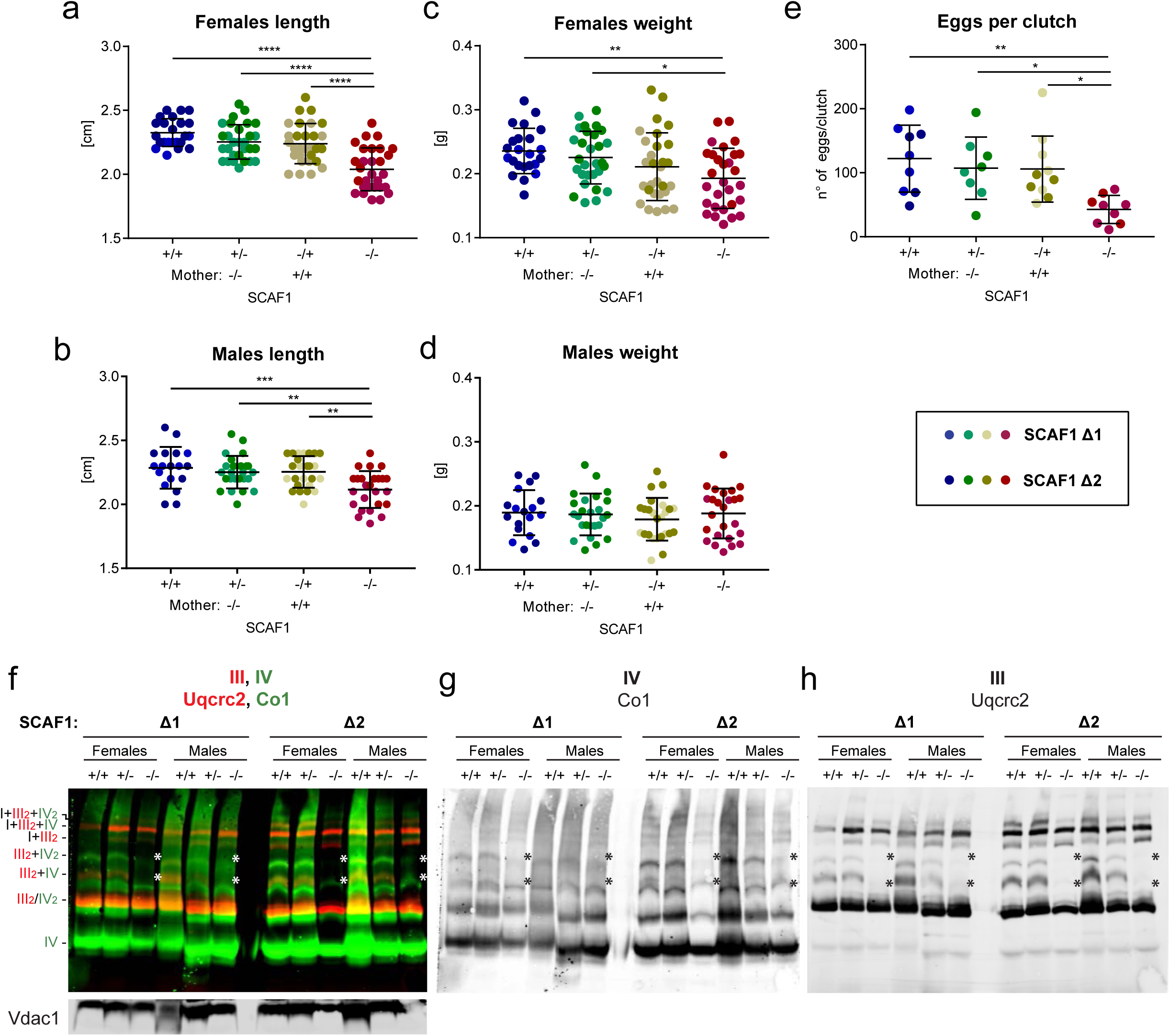
The SCAF1 loss-of-function phenotype is recessive and not maternally contributed. **a-d**, Size of heterozygous offspring from SCAF1^+/+^ and SCAF1^-/-^ females and males. (**a**) Length of F1 females and (**b**) weight (Δ1 +/+ n=7, Δ1 +/- n=16, Δ1 -/+ n=17, Δ1 -/- n=16, Δ2 +/+ n=15, Δ2 +/- n=12, Δ2 -/+ n=11, Δ2 -/- n=12). (**c**) Length of F1 males and (**d**) weight (Δ1 +/+ n=7, Δ1 +/- n=12, Δ1 -/+ n=10, Δ1 -/- n=11, Δ2 +/+ n=10, Δ2 +/- n=12, Δ2 -/+ n=14, Δ2 -/- n=13) **e**, number of eggs per clutch (Δ1 +/+ n=5, Δ1 +/- n=4, Δ1 -/+ n=6, Δ1 -/-, n=6, Δ2 +/+ n=4, Δ2 +/- n=4, Δ2 -/+ n=4, Δ2 -/- n=3). One-way ANOVA. Data are represented as mean ± SD. * p<0.05, ** p<0.01, *** p<0.001, **** p<0.0001. **f-h**, BNGE and immunoblot analysis with the indicated antibodies comparing the impact of SCAF1 deletion in Δ1 and Δ2 fish lines. Asterisks indicate absent bands.

**Figure S8.**
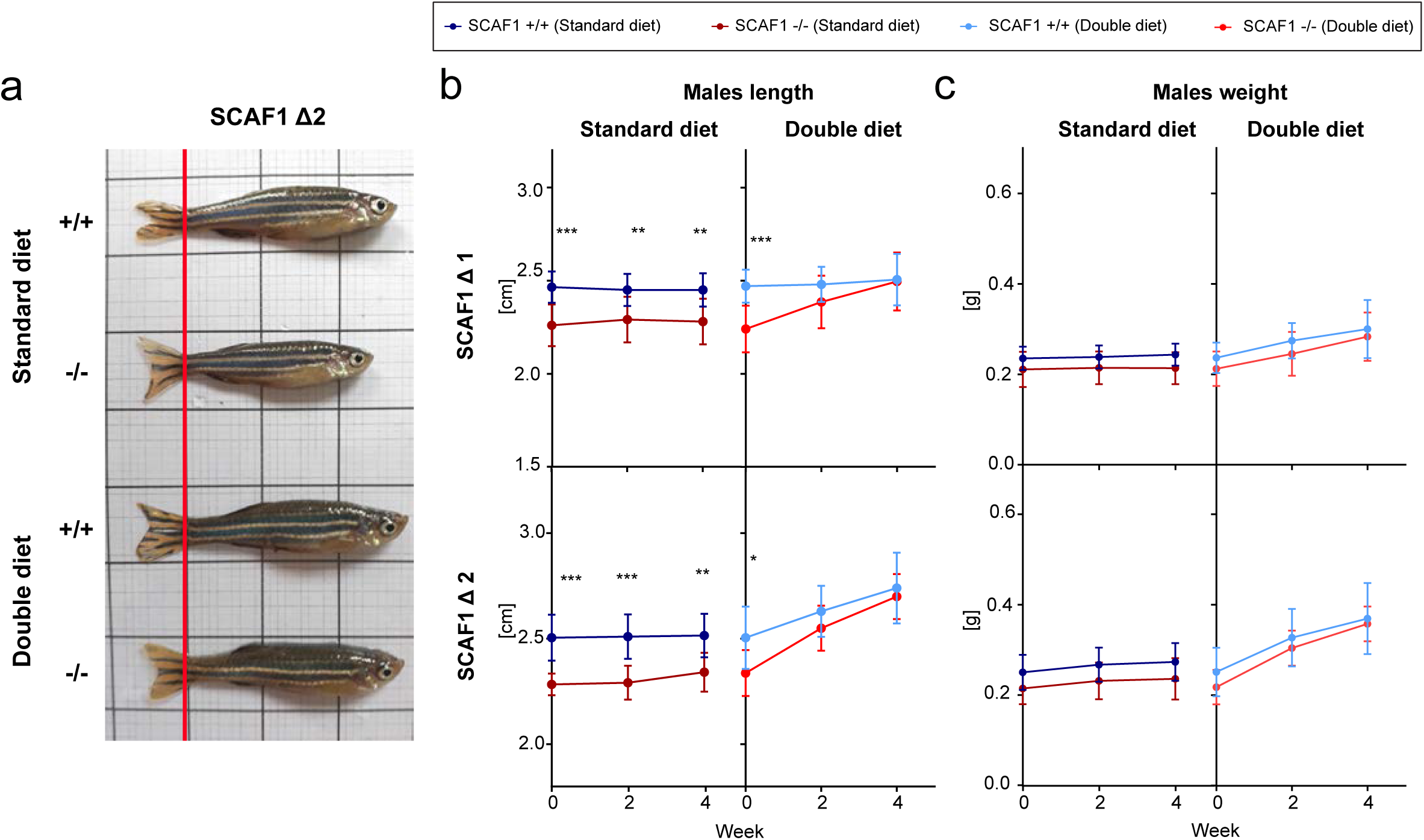
Diet-induced recovery of SCAF1^-/-^ phenotypes in males. **a,** Representative images of SCAF1^-/-^ and SCAF1^+/+^ males fed with the indicated diets. **b,c,** Size of males after the indicated diet. (**b)** Changes in length and (**c**) weight over time (Δ1 +/+ n=10, Δ1 -/- n=10, Δ2 +/+ n=10, Δ2 -/- n=7- 8). Two-way ANOVA. Data are represented as mean ± SD. * p<0.05, ** p<0.01, *** p<0.001.

